# An ancient *cis-*element regulates translation of arginine decarboxylase to control downstream polyamine biosynthesis and stress responses

**DOI:** 10.64898/2026.04.30.721942

**Authors:** Erin Samantha Ritchie, Riëtte Fischer, Edda von Roepenack-Lahaye, Laura Medina-Puche, Ezgi Süheyla Dogan, Xiaofei Yang, Elena Roitsch, Kees Burhman, Tim Michler, Caroline Gutjahr, Martina Ried-Lasi, Yiliang Ding, Chang Liu, Rosa Lozano-Duran, Thomas Lahaye

## Abstract

Polyamines (PAs) are ubiquitous metabolites that, despite their simple structure, profoundly influence plant growth, development, and stress adaptation. Their cellular levels are largely determined by arginine decarboxylase (ADC), a key rate-limiting enzyme in their biosynthesis. We previously identified a ∼50 bp GC-rich sequence in the 5′ untranslated region (UTR) of plant *ADC* genes, termed the *ADC-box*, that is conserved across land plants. Transient reporter assays in tomato, in which ADC upstream regions were decoupled from their native coding sequences and fused to reporter genes, suggested that this element represses translation. However, its function in the native genomic context and its impact on PA homeostasis remain unclear.

Here, we combined CRISPR-Cas9 genome editing, metabolite profiling, enzymatic assays, and RNA structure probing to define *ADC-box* function in tomato and in the seedless land plant *Marchantia polymorpha*, which retains a conserved ∼20 bp core region. Mutation of the *M. polymorpha ADC-box* increased ADC activity and altered PA levels, indicating that the ADC-box functions as a conserved translational repressor. In tomato, disruption of the *ADC-boxes* in *SlADC1* and *SlADC2* increased ADC activity, demonstrating that the *ADC-box* acts as a translational repressor in its native context. These effects were most pronounced under cold stress, when *ADC* transcript levels increase, suggesting that the *ADC-box* buffers stress-induced translation.

Metabolically, *ADC-box* disruption led to agmatine accumulation and alterations in upstream intermediates, while downstream PA pools remained largely unchanged. SHAPE analysis revealed that the tomato ADC-box folds into a three-stem RNA structure, with a central stem representing the major inhibitory module.

*ADC-box* mutants displayed altered plant-microbe interactions, with enhanced resistance to *Pseudomonas syringae* and Tobacco rattle virus, but increased susceptibility to *Ralstonia solanacearum* and Tomato yellow leaf curl virus. Together, these findings establish the *ADC-box* as an evolutionarily conserved *cis*-regulatory element that stabilizes PA homeostasis and modulates plant-microbe interactions.

## Introduction

Polyamines (PA) are low molecular weight metabolites containing at least two amino groups and are essential for almost all organisms ((Michael, 2016)). In plants, PAs play pivotal roles in growth and development (e.g., embryogenesis and flowering), as well as in responses to both abiotic and biotic stresses ((Chen et al., 2019); (Blázquez, 2024)). At a physiological pH, PAs are positively charged and exist either in their free form or bound to cellular components such as nucleic acids, the cell wall, or other metabolites (e.g., hydroxycinnamic acids forming phenolamides) ((Bassard et al., 2010), (Chen et al., 2019)). These diverse interactions complicate the dissection of their precise molecular functions during stress responses.

Regardless, consistent with their role in stress tolerance, numerous studies have generated transgenic plants overexpressing key enzymes of the PA pathway to elevate PA levels. In particular, plants overexpressing a key PA biosynthesis enzyme, arginine decarboxylase (ADC), display enhanced tolerance to salinity, drought, and heavy metal stress ((Pathak et al., 2014)). However, the agricultural use of such transgenic approaches is restricted or poorly accepted in many regions, highlighting the need for alternative strategies to modulate PA levels without introducing transgenes.

In plants, biosynthesis of the major PAs – putrescine, spermidine, and spermine – proceeds via two entry routes ((Chen et al., 2019)). In the primary pathway, arginine is decarboxylated by ADC to produce agmatine, which is subsequently converted into N-carbamoylputrescine and then into putrescine. In the alternative pathway, ornithine can be directly converted into putrescine via ODC. Putrescine is further converted into spermidine and spermine through the sequential addition of aminopropyl groups derived from decarboxylated S-adenosylmethionine (dcSAM). PAs are catabolised by copper-containing amine oxidases (CuAOs) and flavin-containing PA oxidases (PAOs), generating hydrogen peroxide (H₂O₂), a reactive oxygen species that contributes to stress signalling ((Wang et al., 2019)).

A notable feature of the PA network is the prevalence of translational regulation, which likely enables rapid adjustment of enzyme activity and fine-tuning of PA levels in response to changing conditions ((Jimenez-Bremont et al., 2022)). In plants, upstream open reading frames (uORFs) are present in the 5′-untranslated regions (UTRs) of transcripts encoding key PA biosynthetic enzymes, including ADC, PAOs, and *S*-adenosylmethionine decarboxylase, which synthesizes dcSAM, and often mediate regulation in a PA-dependent manner ((Jimenez-Bremont et al., 2022)). In addition, guanine-rich nucleic acid secondary structures such as G-quadruplexes have been identified in PA-related genes, although their functional roles remain largely unexplored ((Yang et al., 2020)).

A distinct translational regulatory cis-element relevant to PA homeostasis was identified in the context of plant–pathogen interactions ((Wu et al., 2019)). The bacterial pathogen *Ralstonia solanacearum* induces the production of truncated *ADC* transcripts in infected tomato (*Solanum lycopersicum*) host tissue, which lack a large portion of the native 5′ UTR and are translated more efficiently. These transcripts are generated by the activity of a type III-secreted effector protein, RipTAL (Ralstonia-injected protein with homology to transcription activator-like effectors), which binds to a 17-nucleotide GC-rich DNA motif – termed the effector binding element (*EBE*) – located approximately 120-140 nucleotides upstream of the start codon of tomato ADCs (SlADC). Binding of the RipTAL induces transcription from an alternative start site located downstream of the *EBE*, producing shorter transcripts with enhanced translational efficiency.

Interestingly, the RipTAL *EBE* is embedded within a larger, highly conserved ∼50-bp GC-rich sequence present not only in *R. solanacearum* host species but broadly across higher plants. This sequence, termed the *ADC-box*, has been shown in reporter assays to repress translation of downstream coding sequences ((Wu et al., 2019)). The absence of the *ADC-box* in RipTAL-induced transcripts likely explains their increased translational efficiency compared to native transcripts containing the full-length 5′ UTR.

In higher plants, the *ADC-box* contains four GC-rich regions (GC1–GC4) that are predicted to form an RNA secondary structure that impedes ribosome scanning and thereby represses translation ((Wu et al., 2019)). A simplified version of this element, comprising only GC2 and GC3, is present in seedless plants such as *Marchantia polymorpha* and is predicted to form a single hairpin structure. However, the function of the *ADC-box* has not been validated in its native genomic context, and the regulatory role of the bryophyte variant remains unknown.

Understanding of how the *ADC-box* functions offers the possibility to modulate endogenous PA levels through targeted sequence modifications. In contrast to transgenic approaches that rely on constitutive overexpression of PA biosynthetic enzymes, manipulation of this cis-regulatory element could enable fine-tuning of ADC expression while preserving native transcriptional regulatory control. Such strategies may also be more compatible with emerging regulatory frameworks and public acceptance.

Given the central role of ADC in PA biosynthesis and the importance of PAs in plant physiology and stress responses, we sought to determine how the *ADC-box* influences ADC protein levels *in planta* and how this affects PA metabolism. To this end, we analysed the *ADC-box* in tomato and the bryophyte *M. polymorpha* using CRISPR–Cas9-mediated mutagenesis. We show that the inhibitory function of the *ADC-box* is conserved from early land plants to higher plants and that it plays a key role in maintaining PA homeostasis. We further examine how disruption of this element affects plant development and responses to abiotic and biotic stresses, and use SHAPE analysis to experimentally define its RNA secondary structure.

## Results

### The ancestral *Marchantia ADC-box* represses translation and maintains polyamine homeostasis

To investigate the functional relevance of the *ADC-box* in the *M. polymorpha ADC* gene, we performed CRISPR–Cas-mediated mutagenesis. Guide RNAs were designed to target either the *ADC-box* (sgRNA1A/B) or a region ∼200 nucleotides upstream (sgRNA2) (Figure 1B; Supplementary Table 1). Because the *MpADC* transcript (MPTK2_4g00150) contains a long 5′UTR, similar to those of seed plants, (>1.5 kb; (Tanizawa et al., 2026)), both target sites are expected to be retained in the mature mRNA. Mutations in the upstream region served as controls, as changes outside the *ADC-box* were not expected to affect translation, allowing *ADC-box*-specific effects to be distinguished from general mutation-induced effects. We identified two lines with upstream control mutations (MpC) and six lines with mutations in distinct regions of the predicted RDRβ hairpin of the *MpADC* transcript (MpM) (Figure 1C, Supplementary Figure 1). The MpM mutants, named by insertion (+) or deletion (−) size, were located predominantly in the GC2-pairing region (MpM-14, MpM-8, MpM+1A), in the loop region between the GC2/GC3 stems (MpM-6), or adjacent to, but outside, the RDRβ hairpin (MpM+1B, MpM+1C). This distribution enabled testing whether the loop is less critical than the paired stem regions, as expected for a structure-dependent regulatory element.

**Figure 1.**
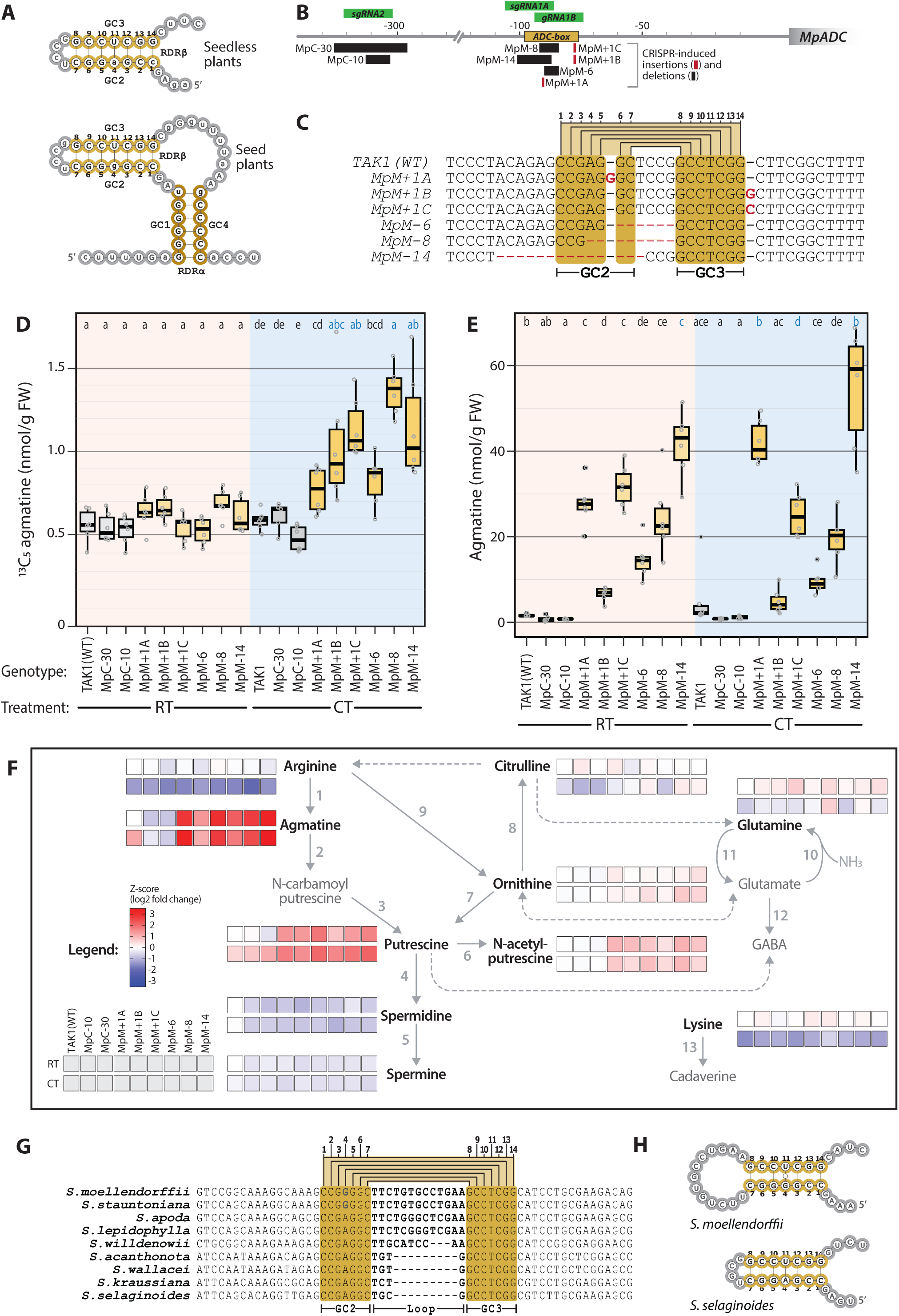
The ancestral *Marchantia* ADC-box represses ADC translation and modulates polyamine homeostasis. (A) Predicted RNA secondary structures of *ADC-box* elements from seedless and seed plants based on previous comparative analyses. The seedless plant *ADC-box* contains the conserved GC2/GC3 regions predicted to form the ancestral RDRβ hairpin, whereas seed plant ADC-boxes additionally contain GC1/GC4 sequences predicted to form the RDRα stem. (B) Schematic of the *Marchantia polymorpha ADC* (MpADC) 5′ untranslated region showing positions of CRISPR/Cas9 guide RNAs targeting either the *ADC-box* (sgRNA1A/B) or an upstream control region (sgRNA2), and locations of recovered mutant alleles. (C) Sequence alignment of *M. polymorpha* WT TAK1 and six independent *ADC-box* mutant alleles. Deleted nucleotides are indicated by red dashes, inserted nucleotides in red font. Conserved GC2 and GC3 pairing regions forming the RDRβ stem are indicated in bold face black font. (D) ADC activity in WT, upstream control mutants (MpC), and *ADC-box* mutants (MpM) maintained at room temperature (RT) or subjected to cold treatment (CT). ADC activity was quantified by LC-MS as formation of ^13^C_5_-agmatine from exogenously supplied ^13^C_6_-arginine. Under CT, most *adc-box* mutants showed significantly increased ADC activity relative to WT and control lines. (E) Endogenous agmatine levels in the same genotypes and treatments shown in (D). Several *ADC-box* mutants accumulated strongly elevated agmatine under both RT and CT conditions. (F) Heat map of polyamine-pathway metabolites quantified from the same extracts and normalized to WT under RT conditions. Values are shown as Z-scores of log2 fold changes. *ADC-box* mutants exhibited strongest increases in metabolites proximal to ADC activity, particularly agmatine and putrescine-related compounds. (G) Alignment of *ADC-box* regions from selected *Selaginella* species. Conserved GC2 and GC3 pairing regions are retained, whereas the intervening loop varies in length among species. (H) Predicted RNA secondary structures of *ADC-box* elements from *Selaginella moellendorCii* and *S. selaginoides*, illustrating tolerated loop expansion while preserving the GC2/GC3 stem core. For boxplots in (D) and (E), center lines indicate medians, boxes indicate interquartile ranges, whiskers indicate 1.5 × interquartile range, and points represent biological replicates. Diierent letters denote statistically significant diierences (ANOVA followed by Tukey’s HSD, P < 0.05).

To assess the impact of *MpADC-box* mutations on translation, we quantified ADC enzymatic activity as a proxy for protein abundance. We used a recently optimized liquid chromatography-mass spectrometry (LC-MS)-based assay ((Ritchie et al., 2025)), in which a stable isotopic variant of the ADC substrate arginine (^13^C_6_ arginine) is added to plant extracts and accumulation of the product ^13^C_5_ agmatine is quantified by LC-MS. The same extracts also permit quantification of additional PA-network metabolites, enabling assessment of broader effects on PA homeostasis. We had previously observed that abiotic stress, including cold treatment (CT), induces *ADC1/2* transcription in tomato and increases ADC activity ((Ritchie et al., 2025)). We therefore reasoned that if *MpADC-box* mutations affect translation, conditions that elevate *MpADC* transcript levels would amplify differences in ADC activity between mutants and wild-type (WT) plants, thereby increasing the sensitivity of the analysis. As CT has previously been shown to induce *MpADC* transcription ((Tan et al., 2023)), the six *MpADC-box* mutants, two control mutants, and the WT TAK1 were either subjected to 16 hours CT or kept at room temperature (RT) control conditions prior to harvesting for LC-MS analysis.

Following CT, all *MpADC-box* mutants except MpM+1A had significantly higher ADC activity (averaging 1.7-2.2-fold) compared to TAK1 and the two control mutants MpC-30 and MpC-10 (Figure 1D). These data suggest that the *ADC-box* is involved in translational regulation of the *MpADC* transcript. Notably, mutants carrying larger deletions within the predicted GC2 paired region (MpM-8 and MpM-14) showed stronger increases in ADC activity than MpM-6, which completely lacks the predicted four-nucleotide loop, yet retains four of the six paired nucleotides forming the adjacent GC2 stem. Together, these data suggest that stem integrity is more important for *ADC-box* function than presence of the predicted loop sequence.

Interestingly, under RT conditions, no significant difference in ADC activity was observed for any *MpADC-box* mutant compared to controls, suggesting that translational control of PA homeostasis by the *ADC-box* is particularly relevant under conditions where *ADC* transcript levels are elevated. By contrast, comparison of PA network metabolites between *MpADC-box* mutants and TAK1 WT controls revealed pronounced metabolic changes not only after CT but also under RT conditions (Figure 1E and 1F), despite increased ADC activity being detectable only after CT (Figure 1D). The strongest effects were observed for agmatine, the immediate product of ADC catalysis. For example, MpM-14 accumulated ∼25-fold higher agmatine levels than TAK1 under RT conditions, despite no significant difference in ADC activity. More moderate, but still substantial increases were observed for putrescine, the downstream decarboxylation product of agmatine, with MpM-14 showing 5.3- and 4.8-fold higher levels than TAK1 under RT and CT conditions, respectively (P < 0.000001; Figure 1F, Supplementary Figure 2, Supplementary Table 2). Increases of lower magnitude were also detected for acetyl-putrescine, a derivative of putrescine, and for ornithine, the precursor of arginine in the broader pathway (Supplementary Figure 2, Supplementary Table 2). Thus, the magnitude of metabolite changes generally declined with increasing biochemical distance from ADC, supporting the conclusion that *ADC-box* mutations primarily enhance ADC-dependent metabolic flux.

The apparent discrepancy between ADC activity and accumulation of agmatine and agmatine-derived PAs likely reflects possibly the different temporal resolution of the two measurements: while the activity assay captures ADC enzyme capacity at the time of harvest, endogenous agmatine and other metabolite levels integrate metabolic flux over time. Thus, *ADC-box* mutations may cause partial basal de-repression of MpADC translation that is sufficient to alter steady-state PA pools at RT, while CT-induced MpADC expression amplifies this effect to a level detectable as increased ADC activity. Collectively, these results show that the ancestral *ADC-box* in *M. polymorpha* not only negatively regulates MpADC levels, but also has a considerable impact on PA network homeostasis.

### *Selaginella* loop variation supports a conserved GC2/GC3 structural core

Previous comparisons of *ADC-box* regions from seed and seedless plants showed that sequence conservation in seedless plants is largely restricted to the GC2/GC3 elements predicted to form the ancestral RDRβ hairpin ((Wu et al., 2019)). One notable exception was *Selaginella moellendorCii*, which contained a nine-nucleotide insertion between the GC2 and GC3 stem regions that was absent from other species analyzed at the time. In light of our mutational data supporting a loop separating the GC2 and GC3 stems, we extended this analysis to additional seedless plants, with emphasis on newly available *Selaginella* sequences. In total, we analyzed 20 *ADC* genes from seedless species, including nine *Selaginella* species. Several *Selaginella* species contained an expanded loop region, whereas others retained the shorter ancestral configuration (Figure 1G and 1H; Supplementary Figure 3). These data indicate that loop expansion arose within the *Selaginella* lineage and that loop length is variable. Together, these results support a stem–loop model in which the paired GC2/GC3 regions form the conserved functional core, whereas the intervening loop is more flexible in length and sequence composition.

### Tomato *ADC-boxes* repress translation in their native genomic context

Having established in *M. polymorpha* that the ancestral RDRβ hairpin functions as a translational repressor controlling PA homeostasis, we next asked whether this conserved structure similarly regulates tomato SlADC1 and SlADC2 levels. Previous work on the tomato *ADC-box* studied GFP reporters driven by *SlADC1/2* 5′UTRs with or without the *ADC-box* in both *in vitro* translation assays and transiently *in planta* ((Wu et al., 2019)), and found that *ADC-box* deletion increased GFP fluorescence by ∼2.5-fold, consistent with *ADC-boxes* acting as translational repressors. However, these reporter assays did not assess *ADC-box* function in its native genomic context and, critically, cannot reveal how the *ADC-box* contributes to maintaining PA homeostasis, because such reporters decouple translation from endogenous ADC expression and metabolism.

### CRISPR-generated tomato ADC-box alleles separate EBE and translational functions

In principle, a previously generated CRISPR/Cas9-edited tomato line could have been used to study *SlADC1* and *SlADC2 ADC-box* function *in planta*. This line was originally designed to disrupt the embedded RipTAL effector-binding elements (*EBEs*) within the *ADC-box* regions of *SlADC1* and *SlADC2*, accordingly designated *Δ1/2-ebe*, and used to confirm RipTAL-dependent activation of host *SlADC1/2* genes ((Wu et al., 2019)).

Re-analysis of the *Δ1/2-ebe* mutations indicated that all bases predicted to form the central RDRβ stem are preserved in both *ADC-boxes* of *SlADC1* and *SlADC2* (termed *SlADC1-box* and *SlADC2-box*), making it likely that their regulatory activity is retained (Figure 2). To better assess *ADC-box* function and its role in PA homeostasis in tomato, we therefore performed a new round of CRISPR-Cas9-mediated to generate lines in which substantial parts, or the entirety of the *SlADC1/2-box* RDRβ stem was removed2. In the T0 generation, we identified one plant carrying a 108 bp deletion that removes the entire *SlADC1-box*, designated *ΔSladc1-box*, and a second plant with a 13 bp deletion within the *SlADC2-box*, designated *ΔSladc2-box* (Figure 2A and 2B, Supplementary Figure 4). As this deletion removed six of the seven nucleotides comprising the GC2 stem sequence, it is expected to impair *SlADC2-box* functionality. T1 progeny were genotyped to identify individuals homozygous for each *ADC-box* mutation and lacking the *Cas9* transgene. Homozygous *Δadc1-box* and *Δadc2-box* plants were then crossed, and subsequent generations were screened to isolate plants homozygous for both mutations, yielding the *Δadc1/2-box* double mutant. In parallel, the previously generated *Δ1/2-ebe* line, which carries a 1 bp insertion in the *SlADC1 EBE* and a 4 bp deletion in the *SlADC2 EBE*, was backcrossed to WT. Genotyping of segregating progeny identified plants carrying either the *SlADC1* mutation (*Δ1-ebe*) or the *SlADC2* mutation (*Δ2-ebe*) (Figure 2A and 2B, Supplementary Figure 4). Together, this generated a set of single mutant lines affecting *SlADC1* (*Δ1-ebe*; *Δadc1-box*) or *SlADC2* (*Δ2-ebe*; *Δadc2-box*), alongside two double-mutant lines affecting both paralogues (*Δ1/2-ebe* and *Δadc1/2-box*).

**Figure 2.**
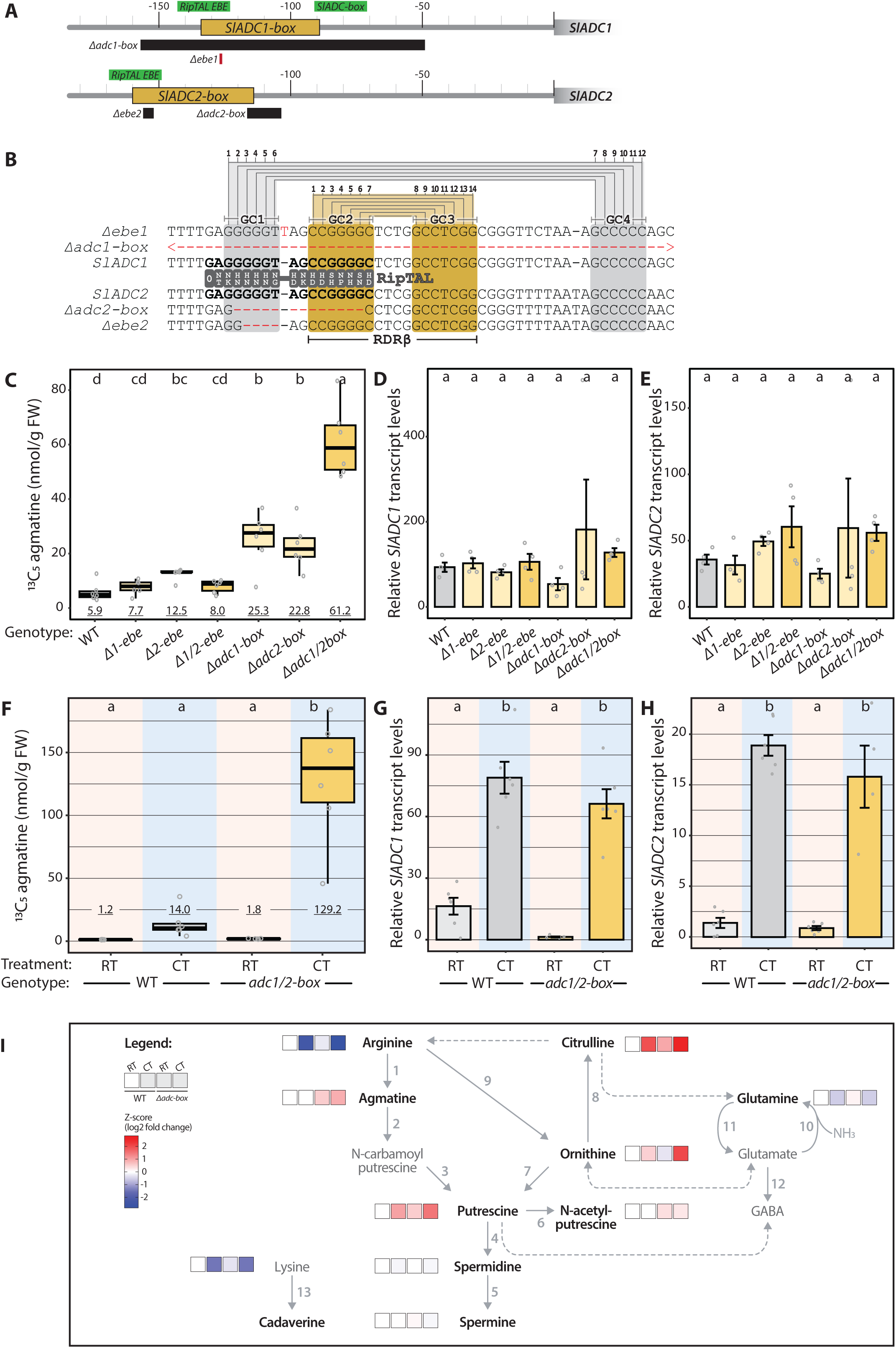
The *ADC-box* represses ADC translation in tomato in its native genomic context and modulates polyamine metabolism. (A) Schematic representation of the *Solanum lycopersicum ADC1* and *ADC2* (*SlADC1/2*) 5′ untranslated regions showing positions of the *ADC-box* (yellow box), embedded sgRNA (green boxes), and CRISPR/Cas9-induced deletions (black boxes) and insertions (red boxes). Image is to scale. (B) Sequence alignments of WT and mutant *SlADC1* and *SlADC2 ADC-box* regions. Conserved GC-rich elements (GC1–GC4) and the central RDRβ pairing region are indicated. Gray boxes with white letters represent the RVDs of a RipTAL protein with nucleotides of the RipTAL target EBE depicted in boldface font. Small *EBE* mutations retain the RDRβ stem, whereas *adc1-box* and *adc2-box* deletions disrupt or remove this structure. (C) ADC activity in WT and mutant lines following cold treatment (CT), quantified as formation of ^13^C_5_-agmatine from ^13^C_6_-arginine. Mutants carrying *ADC-box* deletions show strongly increased ADC activity compared with WT and *EBE* mutants. The average ADC activity is depicted as underlined text. (D, E) Relative transcript levels of *SlADC1* (D) and *SlADC2* (E) in the same samples used for ADC activity measurements. No significant diierences in transcript abundance were detected between genotypes, indicating that ADC-box-mediated regulation acts at the translational level. (F) Comparison of ADC activity in WT and *Δadc1/2-box* plants under room temperature (RT) and CT conditions. The average ADC activity is depicted as underlined text. The *Δadc1/2-box* mutant shows strongly elevated ADC activity relative to WT following CT, whereas diierences are minimal under RT conditions. (G, H) Relative *SlADC1* (G) and *SlADC2* (H) transcript levels under RT and CT conditions. CT induces transcript accumulation in both genotypes without significant diierences between WT and *Δadc1/2*-box plants. (I) Heat map representation of PA-pathway metabolites quantified from the same extracts and normalized to WT controls. Values are shown as Z-scores of log₂ fold changes. Disruption of the ADC-box primarily increases metabolites proximal to ADC activity, particularly agmatine, while downstream polyamines remain comparatively buiered. For boxplots in (C) and (F), center lines indicate medians, boxes indicate interquartile ranges, whiskers indicate 1.5 × interquartile range, and points represent biological replicates. Diierent letters denote statistically significant diierences (ANOVA followed by Tukey’s HSD, P < 0.05). All LC-MS–quantified metabolites were normalized to an internal D₅-tryptophane control.

### Disruption of the RDRβ stem increases ADC activity without altering transcript abundance

To determine how these *SlADC-box* mutations affect translation *in planta*, we again quantified ADC enzymatic activity as a proxy for protein abundance using the LC-MS-based assay. Detached leaves from WT and the six mutant plants explained above were subjected to CT before LC-MS analysis. Mutants included *Δebe* lines carrying mutations in the predicted RipTAL *EBE* while retaining the RDRβ stem *(Δ1-ebe*, *Δ2-ebe*, and *Δ1/2-ebe*), and *Δadc-box* lines carrying larger deletions that removed six of the seven GC2 stem bases of the RDRβ stem or the entire RDRβ stem *(Δadc1-box, Δadc-box, Δadc1/2-box*). Small *EBE*-site mutations (*Δ1-ebe*, *Δ2-ebe*, and *Δ1/2-ebe*) caused little or no increase in ADC activity relative to WT, although *Δ2-ebe* showed a modest but statistically significant elevation (Figure 2C). In contrast, mutants carrying larger *ADC-box* deletions displayed pronounced increases in ADC activity. The *Δadc1-box* and *Δadc2-box* single mutants showed 4.3- and 3.8-fold higher ADC activity than WT, whereas the *Δadc1/2-box* double mutant exhibited a 10.3-fold increase (Figure 2C). These results are consistent with the structural consequences of the mutations: the *Δebe* alleles preserve the central RDRβ stem, whereas the *Δadc-box* alleles disrupt or remove it. The near-additive effect of the double mutant further suggests that the two *SlADC1/2*-boxes act largely independently to repress translation, with little evidence for a higher-order compensatory mechanism that maintains ADC or PA homeostasis under these conditions. Together, these data demonstrate that both *SlADC1/2-boxes* function as translational repressors *in planta* and that repression depends on preservation of the RDRβ stem.

To confirm that the increased ADC activity in the mutants reflects loss of translational repression rather than altered transcript abundance, we quantified *SlADC1* and *SlADC2* transcripts from the same leaf samples as the LC-MS analysis. No significant differences in *SlADC1* or *SlADC2* transcript abundance were detected between any *SlADC-box* mutant and WT plants (Figures 2D and 2E). This supports the conclusion that the *ADC-box* regulates ADC protein abundance at the translational rather than transcriptional level.

### ADC-box repression becomes most apparent under cold-induced transcript accumulation

Because the preceding experiment was performed after CT, which induces *SlADC1/2* transcription and thereby increases the sensitivity for detecting *ADC-box* effects on translation and protein accumulation, we next asked whether the *ADC-box* also influences ADC levels under RT conditions. Leaves from WT plants and the *Δadc1/2-box* mutant, which showed the strongest effect in the initial screen, were therefore maintained at RT or subjected to CT before ADC activity measurements.

Comparing genotypes within each treatment, the *Δadc1/2-box* mutant showed markedly higher ADC activity than WT after CT (9.2-fold), whereas no significant difference between the two genotypes was detected under RT conditions (Figure 2F). Comparing treatments within each genotype, CT caused only a non-significant increase in ADC activity in WT plants, but a strong 71.6-fold increase in the *Δadc1/2-box* mutant.

One possible explanation for this enhanced CT response was that the mutant accumulates higher levels of *SlADC1* and *SlADC2* transcripts. However, CT strongly increased *SlADC1* and *SlADC2* transcript abundance in both genotypes, with no significant differences between WT and *Δadc1/2-box* plants under either condition (Figures 2G and 2H). These data indicate that the pronounced CT-dependent increase in ADC activity of the *Δadc1/2-box* mutant is caused by enhanced translation rather than altered transcript accumulation. Together, this shows that the inhibitory impact of the *SlADC1/2-boxes* becomes particularly evident when *SlADC1/2* transcript levels are elevated.

### ADC-box disruption primarily alters anabolic members of polyamine metabolism

From the same LC-MS extracts used for ADC activity measurements, we quantified ten PAs and related metabolites to assess how increased ADC activity in the *Δadc1/2-box* mutant affects PA homeostasis. Following CT, the *Δadc1/2-box* mutant accumulated 3.7-fold higher agmatine and 9.4-fold higher ornithine levels than WT (P < 0.00001 and P < 0.0009, respectively), whereas arginine, the ADC substrate, was reduced 2.1-fold (P = 0.014; Figure 2I, Supplementary Figure 5, Supplementary Table 3). By contrast, no significant differences were detected for the higher PAs putrescine, spermidine, or spermine, although putrescine showed a reproducible upward trend in the mutant.

Under RT conditions, agmatine, citrulline, and acetyl-putrescine were significantly higher in the *Δadc1/2-box* mutant than in WT plants (P < 0.03, P < 0.0008, and P < 0.008, respectively; Figure 2I, Supplementary Figure 5, Supplementary Table 3). Together, these data confirm that the *ADC-box* functions as a translational regulator of tomato ADC synthesis in its native genomic context and measurably influences PA metabolism.

Notably, the metabolic consequences of *ADC-box* disruption were markedly similar between species. The more substantial *ADC-box* mutations caused increases in agmatine, putrescine, acetyl-putrescine, and ornithine in both *M. polymorpha* and tomato. Yet a key point of difference between the species were the concentrations of the metabolites measured. For example, in *M. polymorpha* quantified agmatine levels ranged from 0 - ∼52 nmol/g FW (Supplementary Table 2), while in tomato this was only 0.5-2.1 nmol/g FW (Supplementary Table 3). On the other hand, arginine land ornithine levels in tomato were up to 350 and 180 nmol/g FW, respectively, while in *M. polymorpha* they only rose to 32 and 1.8 nmol/g FW, respectively. Putrescine levels were also substantially higher in tomato too, up to 195 nmol/g FW (Supplementary Table 3), while only a maximum of 15 nmol/g FW was observed in *M. polymorpha* (Supplementary Table 2). Collectively these results suggest that there are differential metabolic requirements between these species and therefore differential regulatory mechanisms in place to maintain PA pathway flux and nutrient availability.

### SHAPE analysis reveals triple-hairpin structures of tomato ADC-boxes

The *ADC-box* of higher plants has been predicted to form an RNA secondary structure consisting of two stems and a single loop (Figure 1A, Figure 2B; (Wu et al., 2019)). To experimentally validate whether the *ADC-box* forms such a structure *in planta*, we employed Selective 2′-Hydroxyl Acylation analyzed by Primer Extension (SHAPE), a chemical probing approach that quantifies the conformational flexibility of individual nucleotides. In this method, nucleotides are assigned reactivity scores that are subsequently incorporated into computational algorithms to infer RNA secondary structure ((Spitale et al., 2013)). RNA from WT and *Δadc1/2-box* plant tissue was analyzed via SHAPE and a three-stem hairpin structure was deduced for both *SlADC1-* and *SlADC2-boxes* (Figure 3A; Supplementary Figure 6).

**Figure 3.**
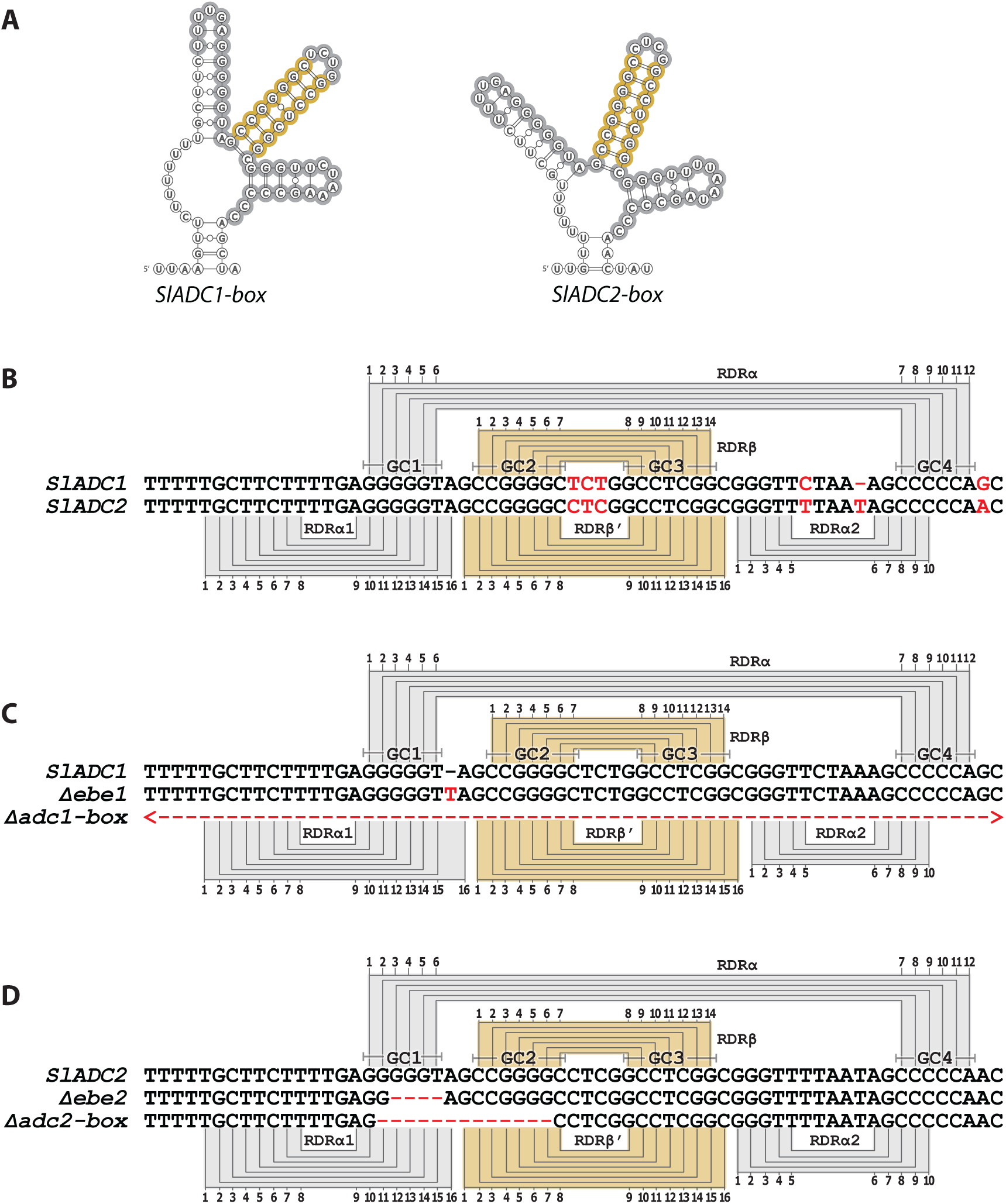
SHAPE analysis reveals triple-hairpin architectures of tomato *ADC-boxes* and identifies RDRβ′ as the principal inhibitory module. (A) SHAPE-guided RNA secondary structures of the *SlADC1-box* and *SlADC2-box* derived from RNA isolated from tomato tissue. Nucleotides engaged in base pairing are indicated. (B) Predicted consequences of representative mutant alleles on ADC-box structure. Small EBE mutations (*Δebe1* and *Δebe2*) largely preserve the overall three-stem architecture, whereas larger ADC-box deletions (*Δadc1-box* and *Δadc2-box*) disrupt the central RDRβ stem region. Corresponding increases in ADC activity measured in Figure 2 are indicated for *Δadc1-box* and *Δadc2-box*. (C) Comparison of the previously predicted two-stem ADC-box model and the SHAPE-derived three-stem model. In the earlier model, GC1 and GC4 form a single RDRα stem and GC2/GC3 form RDRβ. In the SHAPE-supported structure, GC1 and GC4 instead participate in two adjacent stems, designated RDRα1 and RDRα2, whereas the central stem is retained and extended to form RDRβ′. Nucleotide polymorphisms between SlADC1 and SlADC2 occur exclusively in loop regions. (D) Structural interpretation of *SlADC1* mutant alleles. The *Δebe1* mutation leaves the central RDRβ′ stem intact and minimally perturbs adjacent stems, whereas *Δadc1-box* removes the central inhibitory module while leaving flanking structures partially preserved. (E) Structural interpretation of *SlADC2* mutant alleles. The *Δebe2* mutation largely preserves RDRβ′, whereas *Δadc2-box* disrupts the central stem and adjacent paired regions. Together, the SHAPE-derived models explain the mutational phenotypes observed in Figure 2: alleles that preserve RDRβ′ show little eiect on ADC activity, whereas alleles disrupting this stem strongly derepress translation, indicating that RDRβ′ is the dominant structural-functional element of the tomato *ADC-box*.

The experimentally determined paired stems are identical in both *SlADC1-* and *SlADC2-box* sequences, yet remarkably, the few nucleotide polymorphisms between the two *SlADC* paralogs are found exclusively in the predicted loops (Figure 3A, 3B). This is consistent with the hypothesis that there is evolutionary pressure to maintain base pairing within stem regions, whereas loop nucleotides are under weaker structural constraint.

The new SHAPE-derived *ADC-box* structure consists of three stems (Figure 3A, 3B), rather than the two that were previously predicted ((Wu et al., 2019)). The central RDRβ loop is largely conserved between the predicted and experimentally derived models, but now contains eight rather than seven paired nucleotides; we therefore designated this stem RDRβ′. Most GC1 and GC4 nucleotides that were previously predicted to form the RDRα stem instead participate in two independent adjacent stems in the SHAPE-guided model. We designated these new stems RDRα1 and RDRα2, as they contain many nucleotides formerly assigned to RDRα (Figure 3B). Overall, the new three-stem model contains 42 nucleotides engaged in base pairing, compared with 26 nucleotides in the previous two-stem model.

We next asked to what extent this revised three-stem model is consistent with our functional mutagenesis data. In this context, the *Δadc2-box* mutant, which disrupts RDRβ′ but leaves RDRα1 and RDRα2 largely intact (Figure 3C), showed a 3.8-fold increase in ADC activity (Figure 2C), whereas the *Δadc1-box* mutant, in which the complete *SlADC1-box* is deleted (Figure 3D), showed a 4.3-fold increase (Figure 2C). Because these values were not significantly different, the data suggest that loss of RDRβ′ is sufficient to relieve most translational repression, whereas additional removal of RDRα1/2 provides little further effect. Thus, RDRβ′ appears to be the dominant inhibitory structural element.

Although the data indicate that RDRβ′ is the major functional component of the *ADC-box*, comparison of the two *Δebe* mutants suggests that adjacent stems may make smaller contributions. Both *Δ1-ebe* and *Δ2-ebe* retain the RDRβ′ stem and show only minor effects on ADC activity (Figure 2C). However, *Δ2-ebe*, but not *Δ1-ebe*, caused a small yet significant increase in ADC activity. This elevated activity correlates with disruption of four of the eight paired bases of RDRα1, whereas *Δ1-ebe* alters only one paired base of this stem. These observations are therefore consistent with a modest accessory role of RDRα1 in translational repression.

In summary, the experimentally informed SHAPE-derived *ADC-box* data support a model in which RDRβ′ constitutes the principal structural-functional module mediating translational repression of SlADC translation, whereas the adjacent RDRα1/2 stems make smaller supportive contributions.

Tomato *Δadc1/2-box* mutant has no dramatic phenotypic diDerence to WT under non-stressed, salt stressed, or heat stressed conditions.

As PA are involved in many aspects of growth and development ((Chen et al., 2019)) and we have previously shown that tomatoes lacking functional *ADC* genes exhibit severe developmental deformations ((Ritchie et al., 2025)), we asked whether mutations in the *ADC-box* would also cause phenotypic changes in tomatoes. When phenotypic analyses of WT and the double *Δadc1/2-box* mutant tomatoes were conducted under standard greenhouse conditions in three independent experiments, no significant differences were observed in plant height (Figure 4A), shoot number (Figure 4B), adult leaf number (Figure 4C), or flowers (Figures 4D) were detected in the three replicate experiments, nor were there any other apparent physical differences observed (Figure 4E). We therefore concluded that under our greenhouse conditions, the *Δadc1/2-box* and WT plants show no apparent phenotypic differences.

**Figure 4.**
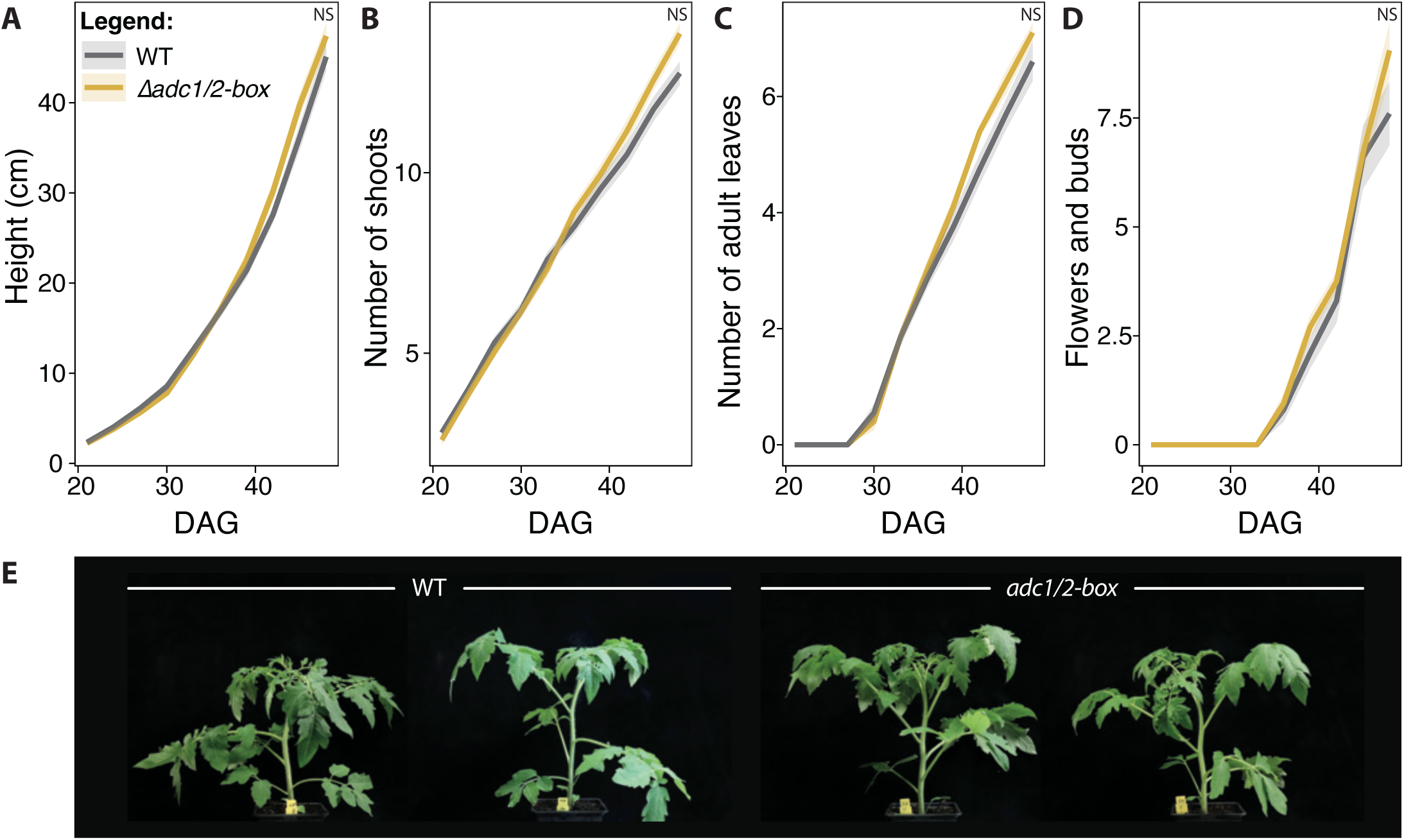
Tomato *Δadc1/2-box* plants show no major developmental phenotype under standard growth conditions. WT (grey) and *Δadc1/2-box* mutant (yellow) tomato plants were grown under greenhouse conditions and phenotyped over time. (A) Plant height, (B) number of shoots, (C) number of fully expanded adult leaves, and (D) number of flowers plus buds were recorded every 3 d from 21 to 48 d after germination (DAG) (n = 20 plants per genotype). (E) Representative plants photographed at 38 DAG. Mixed linear models (A) or generalized mixed linear models (B–D) were used to test genotype eiects across time. No consistent developmental diierences were observed; NS, not significant.

Because no significant differences in ADC activity were detected between WT tomatoes and the *Δadc1/2-box* mutant under mock (RT) conditions (Figure 2F), we hypothesized that a phenotype may become apparent only after *SlADC1/2* transcriptional induction, when the impact of the *Δadc1/2-box* mutation would be amplified. For this, a salt stress was applied as ADCs and PAs have been shown to be protective for tomatoes during salt stress ((Yang et al., 2025)). We also considered that a phenotypic difference between the *Δadc1/2-box* mutant and WT may be observable under conditions where induced ADCs would be detrimental. For this, heat was selected as PA levels and *SlADC1* and *SlADC2* transcript levels have been shown to decrease sharply upon heat stress ((Upadhyay et al., 2020)). However, no statistically significant phenotypic differences were observed between WT and the *Δadc1/2-box* mutant plants, despite clear differences between non-stressed and stressed plants (Supplementary Figure 7). In summary, under both greenhouse and climate chamber conditions, with or without an abiotic stress, the *Δadc1/2-box* mutant showed no apparent phenotype.

### Boosted ADC activity in the *Δadc1/2-box* tomato mutant increases resistance to *Pseudomonas syringae*

PAs have not only been implicated in plant abiotic stress responses, but also in response to biotic stresses ((Jiménez-Bremont et al., 2014); (Gerlin et al., 2021); (Gonzalez et al., 2021); (Blázquez, 2024)). We therefore sought to challenge the tomato *Δadc1/2-box* mutants with a variety of pathogens to test whether altered PA homeostasis affects pathogen susceptibility. *Pseudomonas syringae* pv. *tomato* (*Pto*) is a model bacterial phytopathogen ((Xin and He, 2013)) whose growth negatively correlates with plant PAs in tomato ((Wu et al., 2019)) and *A. thaliana* ((Kim et al., 2013)). We hypothesized that the higher ADC activity in the *Δadc1/2-box* mutants would reduce susceptibility to *Pto* infection. WT and *Δadc1/2-box* tomato leaves were infiltrated with *Pto* or mock and bacterial growth was assessed 2.5 days post infiltration (dpi), when typical *Pto* infection symptoms were observed. As anticipated, *Pto* growth was significantly lower in the *Δadc1/2-box* mutant compared to WT tomatoes (Figure 5A). To understand how ADC activity and PA metabolism are affected by *Pto* infection in both WT and the *Δadc1/2-box* mutant, LC-MS analysis was also conducted on mock- and *Pto*-treated tissue. ADC activity was significantly higher in *Pto*-infected leaves compared to mock in both WT and *Δadc1/2-box* tomatoes (3.9- and 4.8-fold increases, respectively; Figure 5B), most likely due to the induction of *SlADC1* and *SlADC2* transcript abundance following *Pto* infection (Supplementary Figure 8). However, *Pto*-infected *Δadc1/2-box* mutants showed 3.2-fold higher ADC activity than infected WT, indicating that loss of *ADC-box* function elevates ADC protein accumulation. Elevated ADC activity correlated with changes in PA network metabolites, with significantly higher citrulline (4.6-fold), putrescine (2-fold), and ornithine (1.9-fold) levels in the *Δadc1/2-box* mutant compared to WT (ANOVA and Tukey post-hoc test; *P-*values = 0.0074, 0.015, and 0.029, respectively). In addition, arginine (1.8-fold), glutamine (2.2-fold), and spermidine (1.7-fold) levels were significantly lower in the *Δadc1/2-box* mutant (*P-*values = 0.009, 0.04, and 0.0003, respectively). In summary, the *Δadc1/2-box* mutant shows increased resistance to *Pto* compared to WT, which correlates with elevated ADC activity and associated shifts in PA metabolism.

**Figure 5.**
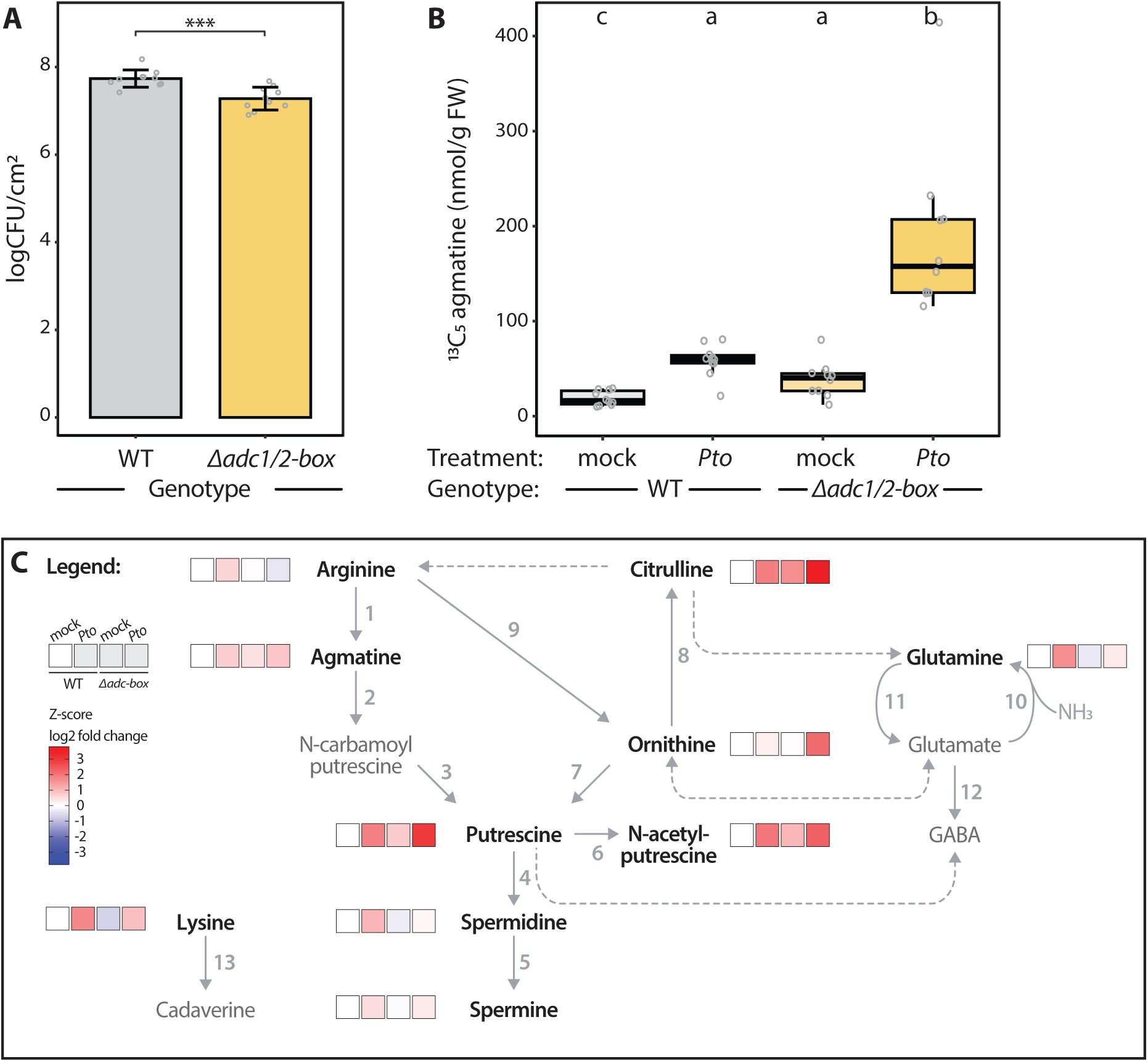
Elevated ADC activity in *Δadc1/2-box* plants enhances resistance to *Pseudomonas syringae pv tomato*. (A) Bacterial growth of *Pseudomonas syringae* pv. *tomato* in WT and *Δadc1/2-box* leaves determined at 2.5 d post infiltration after syringe inoculation. Colony-forming units were quantified from leaf discs. (B) ADC activity in mock- and infected tissue quantified as formation of ^13^C_5_-agmatine from exogenous ^13^C_6_-arginine. (C) Relative *SlADC1* and *SlADC2* transcript abundance in the same samples. (D) Heat map of polyamine-pathway metabolites from infected and mock-treated leaves, normalized to mock WT controls and shown as Z-scores of log_2_ fold change. Infection induced ADC activity in both genotypes, but infected *Δadc1/2-box* tissue accumulated higher ADC activity and altered metabolite profiles relative to WT. Diierent letters indicate statistically significant diierences (ANOVA followed by Tukey’s HSD, P < 0.05).

### Bacterial wilt disease caused by R. solanacearum is accelerated in Δadc1/2-box mutants

The *ADC-box* was originally identified as it contains an effector-binding element (*EBE*) targeted by RipTAL effectors from the *Ralstonia solanacearum* species complex, which, despite extensive sequence divergence across geographically separated phylotypes, retain conserved DNA-binding specificity and activate host *ADC* genes ((Wu et al., 2019); (Gallas et al., 2024)). Given that RipTAL effectors enhance ADC expression during infection, elevated ADC levels would be expected to promote disease. Furthermore, as abiotic stress also induces *SlADC1/2* transcripts (Figure 2G and 2H; (Chen et al., 2019)) and loss of *ADC-box*–mediated repression increases ADC protein accumulation, we asked whether *Δadc1/2-box* tomato mutants show altered susceptibility compared to WT. To investigate this, the disease progression of infected *Δadc1/2-box* and WT tomatoes was assessed following soil drench inoculation with *R. solanacearum*. Because the *ADC-box* contains the RipTAL-binding *EBE*, which is deleted in *Δadc1/2-box* mutants but retained in WT plants, RipTALs would induce *SlADC1/2* expression in WT but not in mutant plants, thereby introducing unequal, host genotype-dependent differences in *ADC* activation during infection. We therefore used a *Δriptal* mutant of the *R. solanacearum* reference strain GMI1000 ((Macho et al., 2010); (Wu et al., 2019)), allowing us to assess the effect of elevated basal ADC activity independently of RipTAL-mediated transcriptional induction. *R. solanacearum* infection proceeds through distinct stages, including root entry, cortical invasion, vascular colonization, and systemic spread, each imposing different constraints on bacterial establishment ((Xue et al., 2020)). To assess whether elevated ADC activity contributes to disease in a stage-specific manner, we performed soil drench infections both with and without prior root wounding. Root wounding facilitates bacterial entry and partially bypasses early infection barriers, thereby shifting the assay towards later stages of infection. The *Δadc1/2-box* mutants were more susceptible to *R. solanacearum* GMI1000 *Δriptal* compared to WT tomatoes both with and without pre-wounding of the roots (Figure 6A and 6B). However, the area under the disease progression curve (AUDPC) was only significantly increased upon root wounding (Figure 6A). These results indicate that elevated ADC levels throughout the *Δadc1/2-box* mutant plant accelerate disease progression with a more pronounced effect when early infection barriers are bypassed.

**Figure 6.**
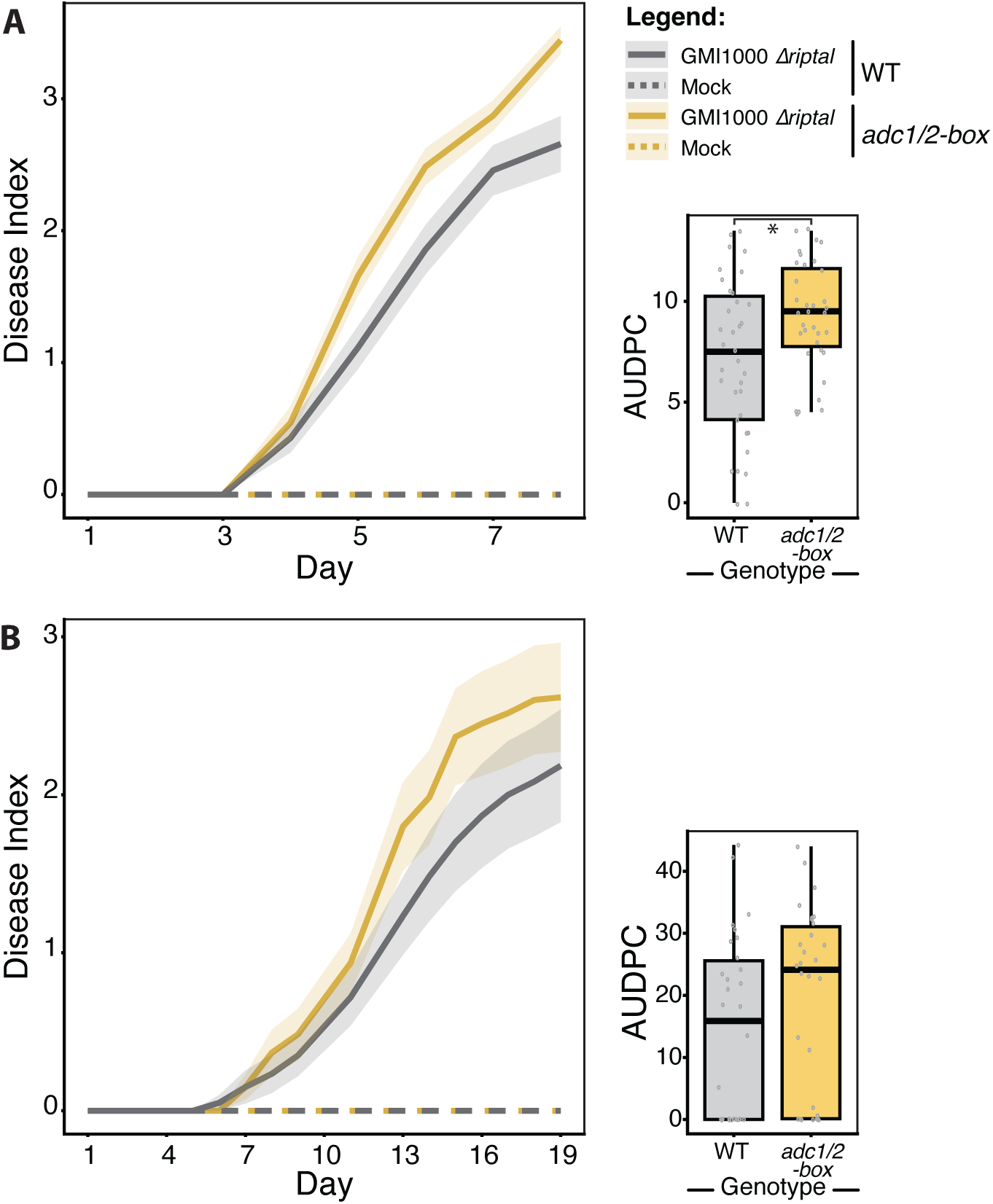
Tomato *Δadc1/2-box* mutants are more susceptible to *Ralstonia solanacearum* infection. WT (grey) and *Δadc1/2-box* mutant (yellow) tomato plants were inoculated with mock (dashed lines) or *Ralstonia solanacearum GMI1000 Δriptal* (solid lines). Disease progression was scored daily using a disease index scale. (A) Root inoculation after pre-wounding. (B) Root inoculation without wounding. Lines indicate means and shaded regions indicate SEM. Insets show area under the disease progress curve (AUDPC) for each genotype. Under wounded inoculation conditions, *Δadc1/2-box* plants developed significantly stronger disease symptoms than WT plants. P < 0.05.

#### The Δadc1/2-box mutant show opposing phenotypes to RNA and DNA viruses

PAs are known to adhere to both DNA and RNA, as well as influence global transcription and translation ((Chen et al., 2019); (Murillo et al., 2025)). We therefore considered how the *ADC-box’s* regulation over the PA network may influence infection by viral pathogens, which are dependent on host transcription and translation. To test this, we analyzed infection by two mechanistically distinct plant viruses: tomato yellow leaf curl virus (TYLCV), a circular single-stranded DNA virus of the genus Begomovirus, and tobacco rattle virus (TRV), a bipartite single-stranded RNA virus ((Lozano-Duran, 2024)). TYLCV accumulation was significantly higher in the *Δadc1/2-box* mutant compared to WT in all three replicate experiments, with an average of 2.6-fold higher levels (Figure 7A). On the other hand, the *Δadc1/2-box* mutants had significantly lower TRV accumulation compared to WT, with an average of 7.5-fold less TRV across the three replicate experiments (Figure 7B). To better understand these contrasting phenotypes, we conducted an ADC activity assay and PA profiling on material from TYLCV- and TRV-treated WT and *Δadc1/2-box* mutant plants. Interestingly, ADC activity was significantly higher in the *Δadc1/2-box* mutant following infection with both TYLCV and TRV compared to WT (Figures 7C and 7D). Similarly, significant increases in agmatine and acetylputrescine in the *Δadc1/2-box* mutants compared to WT were observed during both viral infections, while a significant increase in putrescine levels was only seen in the TYLCV infected *Δadc1/2-box* mutant compared to WT (Figure 7E). Altogether, it seems that mutating the *ADC-boxes* results in opposing phenotypes for TYLCV and TRV infections, despite relatively similar PA and ADC activity profiles being observed during both infections.

**Figure 7.**
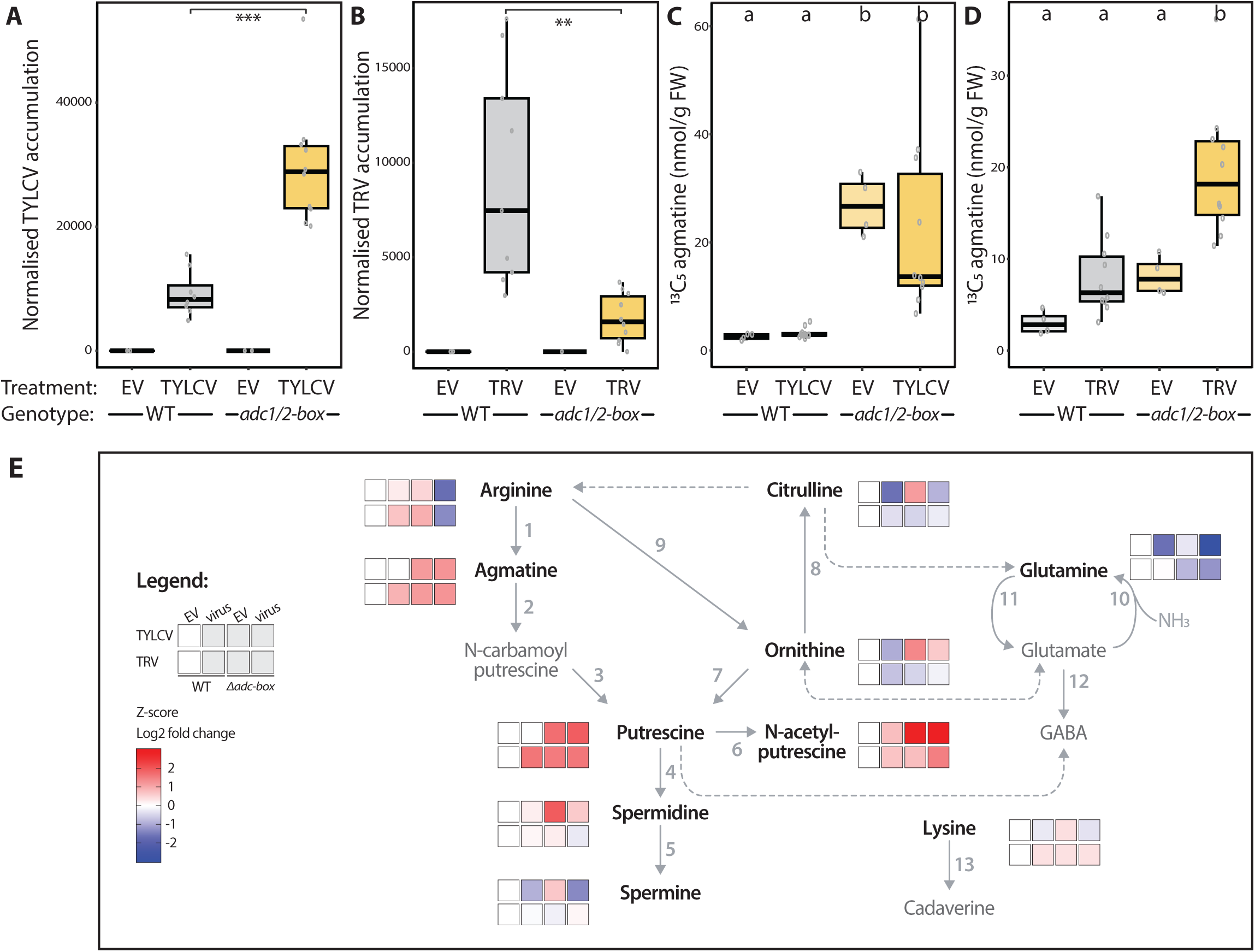
Tomato *Δadc1/2-box* mutants are more susceptible to TYLCV but more resistant to TRV. (A) Accumulation of Tomato yellow leaf curl virus (TYLCV) DNA in WT and *Δadc1/2-box* plants 14 d after inoculation, normalized to *SlActin*. (B) Accumulation of Tobacco rattle virus (TRV) RNA in the same genotypes 14 d after inoculation, normalized to *SlActin*. Empty vector (EV) controls are shown. (C, D) ADC activity in TYLCV- (C) or TRV-infected (D) tissue quantified by LC-MS as ^13^C_5_-agmatine production. (E) Heat map of polyamine-related metabolites from infected samples normalized to EV-infected WT controls and displayed as Z-scores of log_2_ fold change. *Δadc1/2-box* plants supported higher TYLCV accumulation but lower TRV accumulation than WT plants. Asterisks denote significant diierences determined by Student’s t-test; diierent letters indicate significant diierences by ANOVA followed by Tukey’s HSD.

#### The tomato Δadc1/2-box mutant is more resistant to Phytophthora infestans infection

We next investigated how the *Δadc1/2-box* mutant responded to phytopathogen from a different order, such as the oomycete pathogen *Phytophthora infestans* ((Fry, 2008)). First, detached WT and *Δadc1/2-box* mutant leaves were infected with a droplet of *P. infestans* solution, and the resulting lesion size was determined (Figure 8A). The average lesion size on WT plants was 1.7 times greater than those on the *Δadc1/2-box* mutants (Figure 8B). Next, as a more natural infection method, *P. infestans* was sprayed onto the leaf surface and after 5 days the biomass was determined via RTqPCR (Figure 8C). Although lower *P. infestans* biomass was observed in all three replicate experiments, none showed a significant difference (Figure 8D). Together, these results suggest that the *Δadc1/2-box* mutant may be more resistant to *P. infestans*, yet the effect may be too minor too reliably observe.

**Figure 8.**
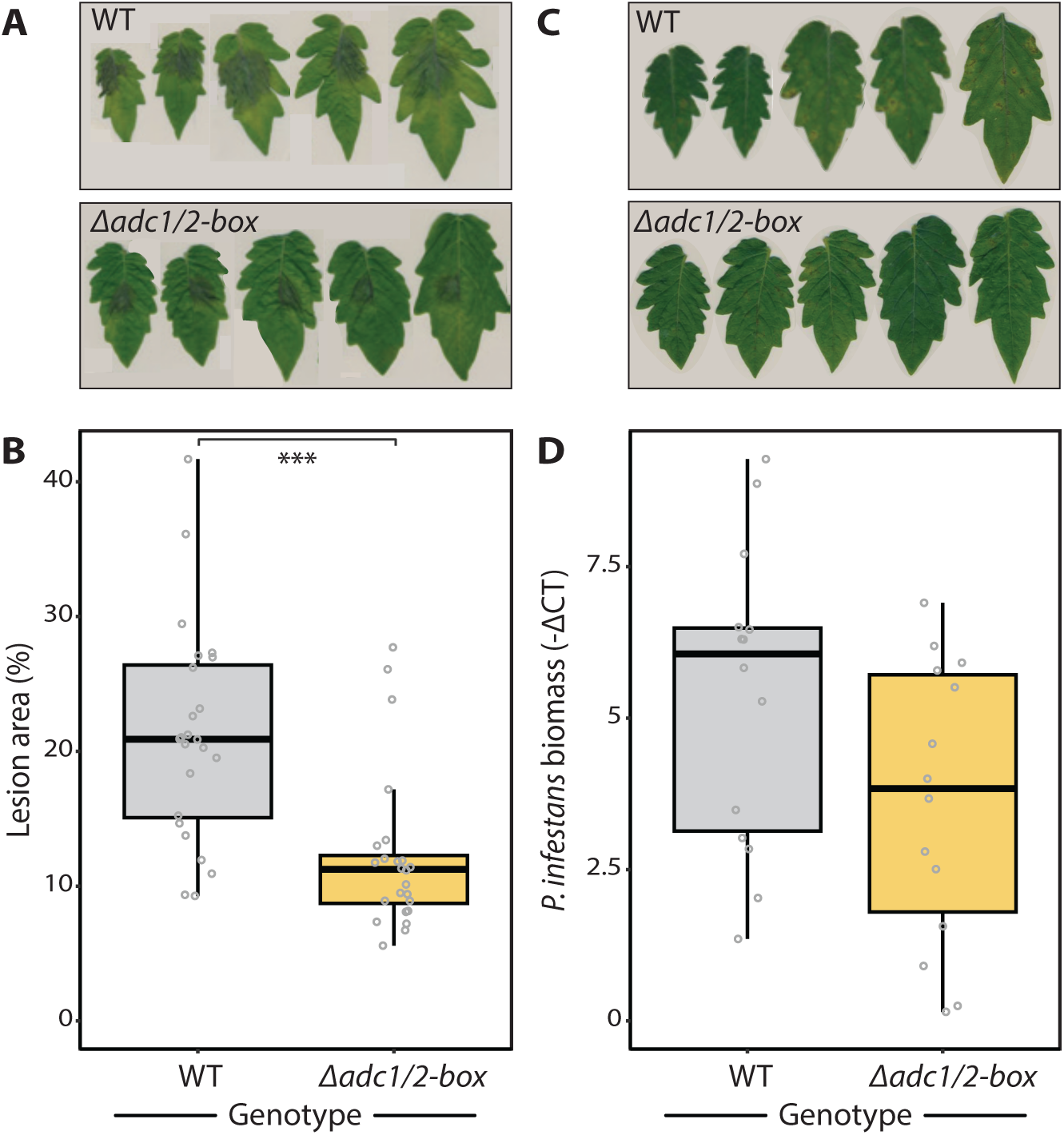
Tomato *Δadc1/2-box* mutants may be more resistant to *Phytophthora infestans*. WT and *Δadc1/2-box* tomato plants were challenged with *Phytophthora infestans* using two infection systems. (A, B) Detached-leaf droplet inoculation. Representative lesions (A) and quantification of lesion area as percentage of total leaf area 7 d after inoculation (B) (n = 24). (C, D) Whole-plant spray inoculation. (C) Representative plants after infection. (D) Pathogen biomass at 5 d after inoculation quantified by qPCR and normalized to the tomato reference gene SlAND (n = 14). *Δadc1/2-box* leaves developed significantly smaller lesions in detached-leaf assays, whereas the reduction in biomass after spray inoculation was not significant. **P < 0.001.

### The *Δadc1/2-box* mutant has lower arbuscular mycorrhiza colonization

Thus far a range of pathogenic phenotypes have been observed on the tomato *Δadc1/2-box* mutants, with increased susceptibility to *R. solanacearum* and TYLCV, but increased resistance to *Pto*, TRV, and possibly *P. infestans* compared to WT. However, we also questioned whether the *ADC-box* may influence PA homeostasis during a non-pathogenic microbial interaction. Therefore, we measured rates of arbuscular mycorrhiza (AM) colonization of the *Δadc1/2-box* tomato mutants compared to WT controls. We observed significantly lower (1.7-fold) AM colonization in the tomato Δ*adc1/2-box* mutants compared to WT, suggesting that the higher endogenous ADC activity and PA levels of this mutant inhibit AM colonization (Figure 9).

**Figure 9.**
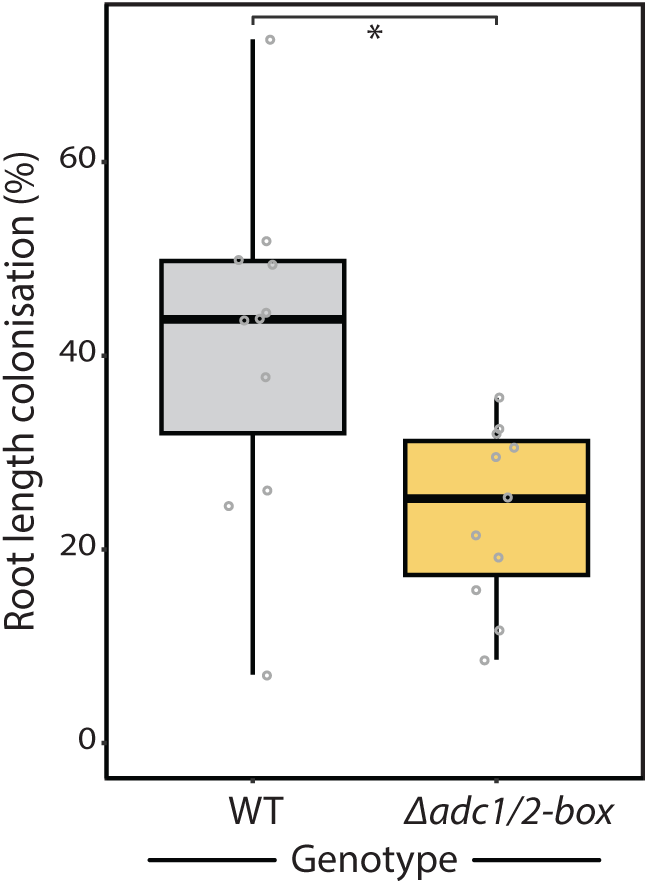
Arbuscular mycorrhizal colonization is reduced in tomato *Δadc1/2*-box roots. Ten-day-old WT and *Δadc1/2-box* tomato seedlings were inoculated with *Rhizophagus irregularis* spores and grown under symbiosis-promoting conditions. After 6 weeks, root systems were harvested, stained, and scored for fungal colonization. Total root length colonization (%) is shown for each genotype (n = 11). *Δadc1/2-box* roots displayed significantly reduced colonization compared with WT roots, indicating that ADC-box-mediated control of PA homeostasis contributes to eiicient arbuscular mycorrhizal symbiosis. P < 0.05.

## Discussion

### The ancestral *ADC-box* in seedless plants is a negative regulatory cis-element of ADC synthesis

The *ADC-box* is an incredibly conserved sequence, with the two central GC-rich regions emerging in early land plants (i.e., bryophytes) and remaining nearly unchanged throughout plant evolution. The GC2 and GC3 regions of seedless plant *ADC-boxes* were previously predicted to form a single RNA hairpin (RDRβ), whereas in higher plants, the acquisition of two additional GC-rich regions (i.e., GC1 and GC4) was proposed to result in the formation of an addition stem, RDRα (Figure 1A; (Wu et al., 2019)).

Here, we investigated the function of the ancestral *ADC-box* found in the bryophyte *M. polymorpha* using CRISPR-Cas9 mutagenesis. ADC activity assays revealed that disruption of the *MpADC-box* led to significantly increased MpADC levels, demonstrating that this ancestral *ADC-box* acts as a negative regulator of downstream translation in its native context (Figure 1D). This translational de-repression was accompanied by substantial metabolic changes: agmatine levels increased by up to ∼25-fold in some mutants, while putrescine levels rose ∼5-fold (Figures 1E and 1F; Supplementary Figure 2). In addition, *M. polymorpha* mutants with more extensive disruption of the *ADC-box* exhibited elevated levels of acetyl putrescine, ornithine, and citrulline, indicating broader perturbation of polyamine metabolism.

Together, these findings demonstrate that even the minimal ancestral *ADC-box*, consisting of only two GC-rich regions, exerts strong regulatory control over ADC abundance and PA homeostasis. Notably, this ∼20-nucleotide RNA element is sufficient to influence not only ADC levels, but also the wider PA metabolic network, highlighting a remarkably compact and effective mechanism of translational regulation.

### *ADC-box* of higher plants also regulates ADC translation and PA homeostasis *in planta* and adopts a triple hairpin structure

Tomato, like most higher plants, has two *ADC* genes, each containing an *ADC-box* with four GC-rich regions previously predicted to form RDRα and RDRβ stem structures. CRISPR-Cas9 mutagenesis of these elements in tomato demonstrated that both *SlADC1-*and *SlADC2-boxes* act as dominant negative regulators of SlADC1 and SlADC2 translation, respectively, *in planta*. The individual tomato *ADC-box* mutants *Δadc1-box* and *Δadc2-box* exhibited ∼4-fold higher ADC activity compared to WT following CT, while the double *Δadc1/2-box* mutant had ∼10-fold higher ADC activity (Figure 2C and 2F). Importantly, these mutants had no significant difference in *SlADC1* or *SlADC2* transcript abundance compared to WT (Figure 2D, 2E, 2G, and 2F), confirming that the *ADC-box* acts at the level of translation, not transcription. As all previous investigations were conducted only using transient reporter assays ((Wu et al., 2019)), these findings provide direct evidence that the *ADC-box* acts as a translational cis-regualtory element in its native genomic context.

Notably, smaller mutations within the *ADC-boxes* (i.e., in the RipTAL *EBE*) had minimal impact on ADC activity, even in the double *Δebe1/2* mutant (Figure 2C). This suggests that the *ADC-box* can tolerate some sequence disruptions, while maintaining functionality, consistent with a structurally mediated mechanism of regulation. To directly assess RNA structure in planta and understand how these mutations influence function, we performed SHAPE analysis on *SlADC1* and *SlADC2* 5’UTR sequences from RNA extracted from tomato tissues. In contrast to the previously predicted two-stem structure (Wu et al., 2019), both ADC-boxes adopted a triple hairpin configuration (Figure 3A). This structure differs from previous predictions in that GC1 and GC4 do not form the RDRα single stem, but instead pair with sequences further 5′ and 3′ to generate two distinct flanking hairpins (Figure 3). In contrast, the central hairpin formed by GC2 and GC3 closely resembles the previously proposed RDRβ, albeit extended by an additional G–C base pair. Given that the GC2 and GC3 sequences are also conserved in the *ADC-box* of seedless plants, such as *M.polymorpha*, it is likely that a similar core hairpin structure is maintained across land plant evolution.

### Function of the *ADC-box* depends on a conserved GC2 and GC3 hairpin

By integrating predicted structural models with CRISPR–Cas9-induced mutations and ADC activity measurements, it becomes evident that the central GC2–GC3 region, corresponding to the RDRβ stem, is critical for *ADC-box* function. Mutations disrupting this region consistently resulted in increased ADC activity. For example, the tomato *Δadc2-box* mutant, in which both GC1 and GC2 are affected, exhibited ∼1.8-fold higher ADC activity than the *Δebe2* mutant, where only GC1 is disrupted (Figure 2C). Similarly, in *M. polymorpha*, mutants MpM_-14 and MpM_-8, which harbour substantial alterations in the GC2 region, showed elevated ADC activity, whereas MpM_-6, with mutations largely confined to the predicted loop region, did not differ significantly from wild type in either ADC activity or agmatine levels (Figure 1).

These functional observations are strongly supported by sequence conservation patterns. The GC2 and GC3 nucleotides are nearly invariant across all examined ADC-box sequences, whereas the flanking regions (GC1 and GC4) are only conserved in higher plants (Wu et al., 2019). In contrast, the loop region of the central hairpin shows considerable variability, with SNPs present even between loop nucleotides of *SlADC1-*and *SlADC2-boxes*, as well as insertions of up to nine loop nucleotides in *Selaginella* species (Figure 1G; Supplementary Figure 3). Together, these data indicating that precise loop sequence is not required for function, while GC2 and GC3 are absolutely required to mediate translational repression of ADCs.

The combination of CRISPR-Cas9 mutagenesis in *M. polymorpha* and tomato, alongside the experimentally derived SHAPE structures, provides strong evidence that the *ADC-box* functions as a regulatory RNA element *in planta*. Given the limited number of plant RNA structural elements that have been experimentally validated, the *ADC-box* represents a compelling example of translational control mediated by RNA structure and suggests that similar regulatory mechanisms may be more widespread in plants than currently appreciated.

### Loss of *ADC-box* function does not have phenotypic consequences under normal or abiotically stressed conditions

PAs are involved in a plethora of growth and developmental roles ((Chen et al., 2019); (Blázquez, 2024)) and numerous studies have demonstrated that perturbation of ADC levels can lead to pronounced phenotypes. The tomato double *adc1/adc2* mutant lacking a functional ADC have severe growth defects and fail to produce flowers ((Wu et al., 2019); (Ritchie et al., 2025)). *A. thaliana* double *adc1/adc2* mutants are embryonically lethal ((Urano et al., 2005)), and even silencing of both *AtADC1* and *AtADC2* results in delayed flowering and stunted plants ((Sánchez-Rangel et al., 2016)). Conversely, *A. thaliana* lines overexpressing ADC are also stunted and have delayed flowering ((Alcázar et al., 2005)).

Given these observations, the substantial increase in ADC activity (∼10-fold) in the tomato *Δadc1/2-box* mutant (Figure 2C and 2F) would be expected to result in a detectable phenotype. Yet, no quantified or observable *Δadc1/2-box* phenotype was found under normal greenhouse conditions (Figure 4). We further tested whether a prolonged stress condition (i.e., salt or heat), which transcriptionally activate *SlADC1/2*, might reveal latent phenotypes, but no differences were detected (Supplementary Figure 7).

These findings suggest that disruption of the *ADC-box*, and the subsequent increase in ADC activity, does not phenocopy constitutive ADC overexpression, likely due to the preservation of transcriptional and possibly proteomic regulation, as well as compensatory control within the polyamine metabolic network. We therefore propose that the absence of a phenotype reflects robust homeostatic buffering, whereby long-term readjustment of PA levels is achieved through alternative regulatory mechanisms that maintain PA stability despite elevated ADC activity.

### *ADC-box*-mediated regulation shapes niche-specific outcomes during plant-pathogen interactions

ADCs and PAs are well-known players in plant–microbe interactions ((Jiménez-Bremont et al., 2014); (Gerlin et al., 2021); (Gonzalez et al., 2021); (Blázquez, 2024)). During pathogen attack, PA levels are typically induced, often via transcriptional upregulation of *ADCs*. This induction of PA levels contributes to defense through multiple mechanisms, including ROS production via PA catabolism and reinforcement of cell walls through phenolamide biosynthesis. However, interpretation of PA function in plant–microbe interactions is complicated by the fact that many pathogens also synthesize and, in some cases, secrete substantial amounts of PAs during infection ((Lowe-Power et al., 2018); (Vilas et al., 2018); (Gerlin et al., 2021)).

Here, we investigated the role of *ADC-box*-mediated regulation over ADC synthesis in the context of plant-microbe interactions. We challenged the tomato *Δadc1/2-box* mutant with a diverse panel of pathogens, which revealed contrasting phenotypes: enhanced resistance to *Pto* (Figure 5), TRV (Figure 7), and *P. infestans* (Figure 8), but increased susceptibility to *R. solanacearum* (Figure 6) and TYLCV (Figure 7). The increased resistance to *Pto* is consistent with previous reports showing a negative correlation between ADC levels and bacterial growth in both tomato and *A. thaliana* ((Kim et al., 2013); (Wu et al., 2019); (Liu et al., 2020)). Moreover, induction of *ADC* transcription and PA accumulation has been reported during TRV infection (Fernandez-Calvino et al., 2014), and enhanced PA catabolism has been shown to restrict *P. infestans* infection (Yang *et al*., 2022), supporting a role for PAs in limiting these pathogens.

A plausible explanation for the increased resistance to Pto, TRV, and *P. infestans* is that elevated ADC activity in the *Δadc1/2-box* mutant enhances PA turnover and associated ROS production, thereby restricting pathogen growth during early, typically biotrophic stages of infection. In parallel, ADCs and PAs are closely linked to hormone signaling networks ((Gerlin et al., 2021)), and altered ADC activity may reshape salicylic acid (SA), jasmonic acid, and ethylene crosstalk. For example, ADC activity has been implicated in SA-dependent defense and systemic acquired resistance in *A*. *thaliana* ((Liu et al., 2020)), raising the possibility that increased ADC levels promote SA-associated defenses in the *Δadc1/2-box* mutant. However, this cannot fully explain the observed phenotypes, as TYLCV – whose accumulation is suppressed by SA in tomato ((Li et al., 2019)) – proliferated to higher levels in the *Δadc1/2-box* mutant.

We therefore propose that the contrasting resistance and susceptibility phenotypes observed in the *Δadc1/2-box* mutant compared to WT are not solely determined by defense signaling, but also by pathogen lifestyle and the specific host niche they exploit. In particular, both *R. solanacearum* and TYLCV, which showed enhanced proliferation in the *Δadc1/2-box* mutant, are vascular-associated pathogens. Elevated ADC activity and altered PA metabolism may modify the metabolic or structural properties of vascular tissues, creating a more permissive environment for these pathogens. In contrast, pathogens that rely on early biotrophic colonization may be more sensitive to enhanced ROS-associated defenses.

Collectively, these findings indicate that *ADC-box*-mediated regulation of PA homeostasis differentially shapes plant-microbe interactions, with outcomes determined by the interplay between defense activation, metabolic state, and pathogen niche. To our knowledge, this study provides the first direct genetic evidence for a role of ADCs during infection with TRV, TYLCV, and *P. infestans*. Further work will be required to disentangle the relative contributions of *ADC-box*-dependent changes in metabolism, hormone signaling, and cellular environment to the observed pathogen-specific phenotypes.

### The *ADC-box* is important for establishment of symbiosis with arbuscular mycorrhizal fungi

In addition to pathogenic interactions, we examined the symbiotic association with the arbuscular mycorrhizal fungus *Rhizophagus irregularis* in WT and *Δadc1/2-box* tomato plants. Notably, the *Δadc1/2-box* mutant exhibited a lower level of root colonization compared to WT (Figure 9), suggesting that elevated ADC activity negatively impacts the establishment or maintenance of this symbiosis.

This observation contrasts with previous studies showing that exogenous application of PAs can enhance arbuscular mycorrhizal colonization ((Jiménez-Bremont et al., 2014); (Sharma et al., 2021)), and that symbiosis itself is associated with increased levels of arginine, agmatine, ornithine, and putrescine, as well as elevated ADC activity ((Zou et al., 2020)). However, these studies largely reflect transient or externally supplied increases in PAs, whereas disruption of the *ADC-box* leads to sustained, endogenous elevation of ADC activity and broader metabolic reprogramming. Under the nutrient-limiting conditions required to establish mycorrhizal symbiosis, such systemic changes may alter the plant’s physiological state, making it more unfavourable for symbiotic establishment. For example, the *Δadc1/2-box* mutant may exhibit altered nutrient status or signalling, shifting the balance between defence and symbiosis. This could involve changes in nitrogen allocation, carbon–nutrient exchange, or hormone signalling pathways known to regulate mycorrhizal interactions. In particular, given the metabolic link between PA and ethylene biosynthesis, and the role of ethylene as a negative regulator of arbuscular mycorrhizal colonization, altered hormonal balance in the *Δadc1/2-box* mutant may have contributed to reduced colonization.

Together, our findings show that the *ADC-box* plays an important role in during plant microbe interactions. A clear mechanistic understanding as to why the *Δadc1/2-box* mutant was more resistant to *Pto*, TRV, and *P. infestans*, but more susceptible to *R. solanacearum* and TYLVC is still lacking. We propose the role of PAs may not be dependent on pathogen lifestyle, a notion that was also suggested by (Jiménez-Bremont et al., 2014), but rather due to the niche where the pathogen resides (i.e., vasculature).

## Conclusion

In conclusion, this work establishes the *ADC-box* as a highly conserved and functionally significant cis-regulatory RNA element that modulates ADC translation across land plant evolution. From its minimal ancestral form in bryophytes to the more complex architecture in higher plants, the *ADC-box* retains a remarkably conserved core sequence, underscoring strong evolutionary constraint on its function. Most notably, our structural analyses reveal that the higher plant *ADC-box* adopts an unexpected triple hairpin configuration *in planta*, redefining previous models and highlighting the importance of RNA secondary structure in translational control. The strict conservation of the central GC2–GC3 region, forming the RDRβ stem, and its essential role in regulatory function further point to a shared mechanistic basis maintained over hundreds of millions of years.

Despite substantially elevated ADC activity, the tomato *Δadc1/2-box* mutant does not exhibit a detectable developmental phenotype, suggesting robust buffering of PA homeostasis under standard conditions; however, its altered responses across a range of plant-microbe interactions reveal a context-dependent functional importance of *ADC-box*-mediated regulation.

Beyond its evolutionary significance, the *ADC-box* represents a rare, experimentally validated example of a plant RNA structural element with direct regulatory function, suggesting that similar mechanisms may be more widespread than currently appreciated. Together, these findings position the *ADC-box* as a unique and compelling model for understanding RNA-mediated translational regulation and its integration into plant metabolic and physiological networks.

## Materials and Methods

### Plant growth conditions

Tomato (Moneymaker) plants were grown at 24 ± 2°C, 30-50% humidity in soil. *M. polymorpha* (TAK1) plants were grown at 18 ± 2°C on ½ B5 media (no added sucrose) as per (Sugano et al., 2014). All plants grown in a 16/8-hour light/dark photoperiod.

### CRISPR-Cas9 mutagenesis

Constructs for CRISPR-Cas9 mutagenesis of tomato *ADC-boxes* (*Solyc10g054440.2* and *Solyc01g110440.4*) were made as described by (Jacobs et al., 2015), using the CRISPRdirect online tool (Naito et al., 2015) ensuring no predicted oh targets with the 20-nucleotide gRNA. These constructs were stably transformed described by (Wittmann et al., 2016). CRISPR-Cas9 mutagenesis of *M. polymorpha ADC-box* (*Mp4g00150.1)* was performed as described ((Sugano et al., 2014)).

For identification and genotyping of mutants, the targeted region was amplified from extracted genomic DNA and Sanger sequenced (Supplementary Table 4).

### Cold treatment

CT of tomatoes was performed as described by (Ritchie et al., 2025). *M. polymorpha* were grown from gemma on fresh ½ B5 plates for 3 weeks before either being CT at ∼6°C, in darkness, for 16 hours (overnight) or kept at RT in the dark. LC-MS samples were taken without allowing the plants to warm.

### ADC activity and PA profiling via LC-MS

Quantification of ADC activity and PA metabolites was performed as described by (Ritchie et al., 2025). *M. polymorpha* samples were treated in the same manner, harvesting 50 mg FW tissue from one independent plant per biological replicate.

### RT-qPCR of tomato *ADC1/2*

Measurement of *SlADC1/2* transcript abundance (i.e., RNA extraction and RT-qPCR) was conducted as previously described ((Ritchie et al., 2025), (Wu et al., 2019)) using primers shown in (Supplementary Table 4).

### Phenotyping tomatoes

Tomato phenotyping was conducted according to (Ritchie et al., 2025). For salt stress, tomato seedlings were grown in 7x7x8 cm pots before watering with 125 mM NaCl or water every 3 days. Heat stress experiments were conducted with a 16hr/8hr light/dark cycle, 50% humidity, and either constant 24 °C, or with a daytime heat stress up to 32 °C for 14 hours, with 1 hour to increase or decrease the temperature to 24 °C overnight.

### *Pseudomonas syringae* infection

*Pseudomonas syringae pv. Tomato DC3000* containing an empty pBBR1MC5 vector (EV; (Kovach et al., 1995)) with gentamycin resistance was streaked onto King’s B agar and grown for 2 days at 28 °C. Cells were resuspended in 10 mM MgCl_2_ to an OD_600_ 0.00002 before infiltration into the abaxial leaf surface of ∼4-week-old tomatoes that were then kept in ∼100% humidity. After 2.5 days, 0.5 cm^2^ of tissue was taken and the number of colony forming units (CFU) per cm^2^ was determined.

### Tomato soil drench with *R. solanacearum*

*Ralstonia solanacearum* GMI1000Δ*riptal* (also known as GMI1000Δ*brg11* (Macho et al., 2010)) were grown on CPG agar (1 % peptone, 0.1% casamino acids, 0.5% glucose, 0.005% tetrazolium chloride, 1.8% agar) with gentamycin (10 µg/mL) at 28 °C for 48 hours. A single colony was used to inoculate liquid CP medium ((1 % peptone, 0.1% casamino acids), grown overnight at 28 °C, 180 rpm, before centrifugation (4000 rpm, 10 min), washed, and resuspended in MilliQ water and adjusted to an OD₆₀₀ of 0.01.

Tomatoes (3–4 weeks old, grown in 7 × 7 × 8 cm pots) were not watered for 24 hours prior to infection. Each plant was inoculated by soil drenching with 50 mL of bacterial suspension or MilliQ water as mock. Soil moisture was maintained to prevent drying for several days post-inoculation. Viability of inoculum was confirmed by re-streaking onto CPG agar containing gentamicin and incubating at 28 °C for 48 h. Disease progression was scored every day according to (Morcillo et al., 2020), whereby 0 = no wilting; 1 = ≤24% leaf area wilted; 2 = 25–49%; 3 = 50–74%; 4 = 75–100%.

### Viral infections

Systemic viral infections were performed as described in (Medina-Puche et al., 2020). In brief, suspension of *Agrobacterium tumefaciens* carrying the TYLCV infectious clone (AJ489258; NCBI:txid220938) recombined into pGWB501 ((Nakagawa et al., 2007)), pTRV1, or pTRV2 ((Liu et al., 2002)) were syringe-inoculated in the stem of two-week-old tomato plants. The second youngest apical leaves were harvested at 14 dpi to detect viral accumulation.

### Determination of viral accumulation

For TYLCV infections, total DNA was extracted following the CTAB method (Murray and Thompson, 1980). Viral DNA accumulation was quantified by qPCR with primers to amplify TYLCV Rep (Wang et al., 2017). As internal reference for DNA detection, the *25S ribosomal DNA interspacer* (*ITS*) was used (Mason et al., 2008).

For TRV infections, RNA was extracted following the citrate-citric acid RNA isolation method (Onate-Sanchez and Vicente-Carbajosa, 2008). DNAse I (Thermo Scientific) treated RNA (500 ng) was used for cDNA synthesis using PrimeScript RT MasterMix (Takara) following manufacturer’s instructions. Viral RNA accumulation was quantified by RT-qPCR with primers to amplify TRV and *SlACTIN* as an internal reference for cDNA detection. BioRad CFX384 real-time system with PowerTrack SYBR Green Mastermix (Thermo Scientific) was used for qPCR, with the following program: 95°C (2 min), and 40 cycles of 95 °C (15 sec) and 60 °C (1 min).

### *Phytophthora infestans* infection methods and quantifications: *P. infestans* and tomato culturing

Tomato seeds were germinated directly on soil, then re-potted after one week and cultivated under long-day conditions (16 h light/8 h darkness, 24 °C/24 °C, 150 µmol/m²/s, 50 % humidity).

The *Phytophthora infestans* isolate CRA 208m2 ((Si-Ammour et al., 2003)), kindly provided by Prof. Dr. Sabine Rosahl, was maintained on oat bean sucrose agar medium (34 g/L white bean flour, 17 g/L oat flour, 8.5 g/L sucrose, 15 g/L agar) at 17 °C in darkness. Sporangial suspensions were prepared from 14-day-old cultures by adding sterile water and incubating in the dark for 4 h. Suspensions were filtered (Miracloth, pore size 22-25 µm; Merck) to separate spores from unopened sporangia and mycelium debris. Spore concentration was determined using a LUNA™ Automated Cell Counter (Logos Biosystems) and adjusted to 1 x 10^5^ spores/mL.

### Analysis of Phytophthora infestans abundance

At the stage of 5–6 fully developed compound leaves, leaves at the developmental stage 3 were spray-inoculated on the abaxial side with the *P. infestans* spore suspension. Plant trays were sealed to maintain high humidity. At 5 dpi, nine leaf discs (9 mm diameter) per leaf triplet were collected, pooled, and snap-frozen in liquid nitrogen. Genomic DNA was extracted (GenElute™ Plant Genomic DNA Miniprep Kit; Sigma-Aldrich) according to the manufacturer’s instructions and concentration quantified (Agilent BioTek Epoch Microplate Spectrophotometer; Agilent Technologies).

Pathogen biomass was quantified by RT-qPCR targeting the *P. infestans*-specific genomic repeat sequence PiO8 ((Judelson and Randall, 1998)); tomato SAND gene (*Solyc03g115810*) used as an internal reference. RT-qPCR was performed using PowerUp™ SYBR™ Green Master Mix (ThermoFisher Scientific) and a QuantStudio 5 real-time PCR system (Applied Biosystems) using the following program: 95 °C (3 min), followed by 40 cycles at 95 °C (10 s) and 60 °C (30 s).

### Analysis of lesion size

At the stage of 5–6 fully developed compound leaves, leaves at the developmental stages 3 and 4 were detached and placed on moist paper towels. A 10 µL droplet of *P. infestans* spore suspension was applied to the abaxial side of each leaf, followed by incubation under 100% relative humidity. Leaves were scanned at 7 dpi for lesion assessment. Total leaf area and lesion area were measured from the adaxial side using ImageJ software. Lesion size was normalized to the total leaf area.

### Assessment of Arbuscular mycorrhizal colonization

Tomato seeds were surface sterilized with 1% NaClO and 0.1% SDS, washed with MilliQ water for 5 times, germinated on 0.8% Plant Agar (Duchefa, Netherlands) and kept in darkness (1 week), before put to light. Arbuscular Mycorrhiza experiments were performed in open pots filled with quartz sand/vermiculite (1:2) mixture. Per pot 2 seedlings (∼10 days old) were transplanted with 500 *R. irregularis* DAOM 197198 spores (type C, Agronutrion, Toulouse, France) per plant. Each week, plants were watered twice with B&D medium (50 µM phosphate and 7.5 mM nitrogen) and once with deionized water ((Torabi et al., 2021)). Plants were grown in long-day photoperiod (16h light at 23 °C/8h dark at 21 °C, at 200 µmol/m^2^/s light intensity) for the first 5 weeks then transferred to a high light chamber (12h light at 26 °C /12h dark at 20 °C, programmed at 500 µmol/m^2^/s light intensity). After 6 weeks, roots were harvested and cleared in 10% KOH at 95 °C, 10 minutes, then washed in 10% Acetic acid, and stained in 5% Black ink (Pelikan 4001, *Brilliant Black*) at 95 °C for 5 minutes. After staining, roots were washed and stored in 5% Acetic acid. Root length colonization (RLC) was assessed using a Zeiss stemi 305 stereoscope using a grid-intersect method ((Giovannetti and Mosse, 1980)).

### Data Analysis and Visualization

Data and statistical analyses were conducted using Microsoft Excel and R; all R code is available on github. Figures were assembled using Adobe Illustrator.

## Acknowledgments

We gratefully acknowledge funding from the Deutsche Forschungsgemeinschaft (DFG) of LA 1338/15-1 and TRR356 (project number: 491090170, subproject B03). Metabolite analytics in ZMBP central facilities unit was funded by the DFG (Project number 442641014). We thank Haseong Kim for providing the Pseudomonas syringae EV strain, and Sabine Rosahl for support and resources related to *Phytophthora* work. We are grateful to Natalie Faiss for assistance with *Marchantia polymorpha*, and to Dousheng Wu for generating the *Δebe1/2* mutant cross to WT. We also thank Dennis Perrett for valuable advice on statistical analyses. We acknowledge the excellent support of the greenhouse staff and the plant transformation unit. Finally, we thank all members of the Lahaye laboratory for helpful discussions and critical reading of the manuscript.

---

**Supplementary Figure 1.**
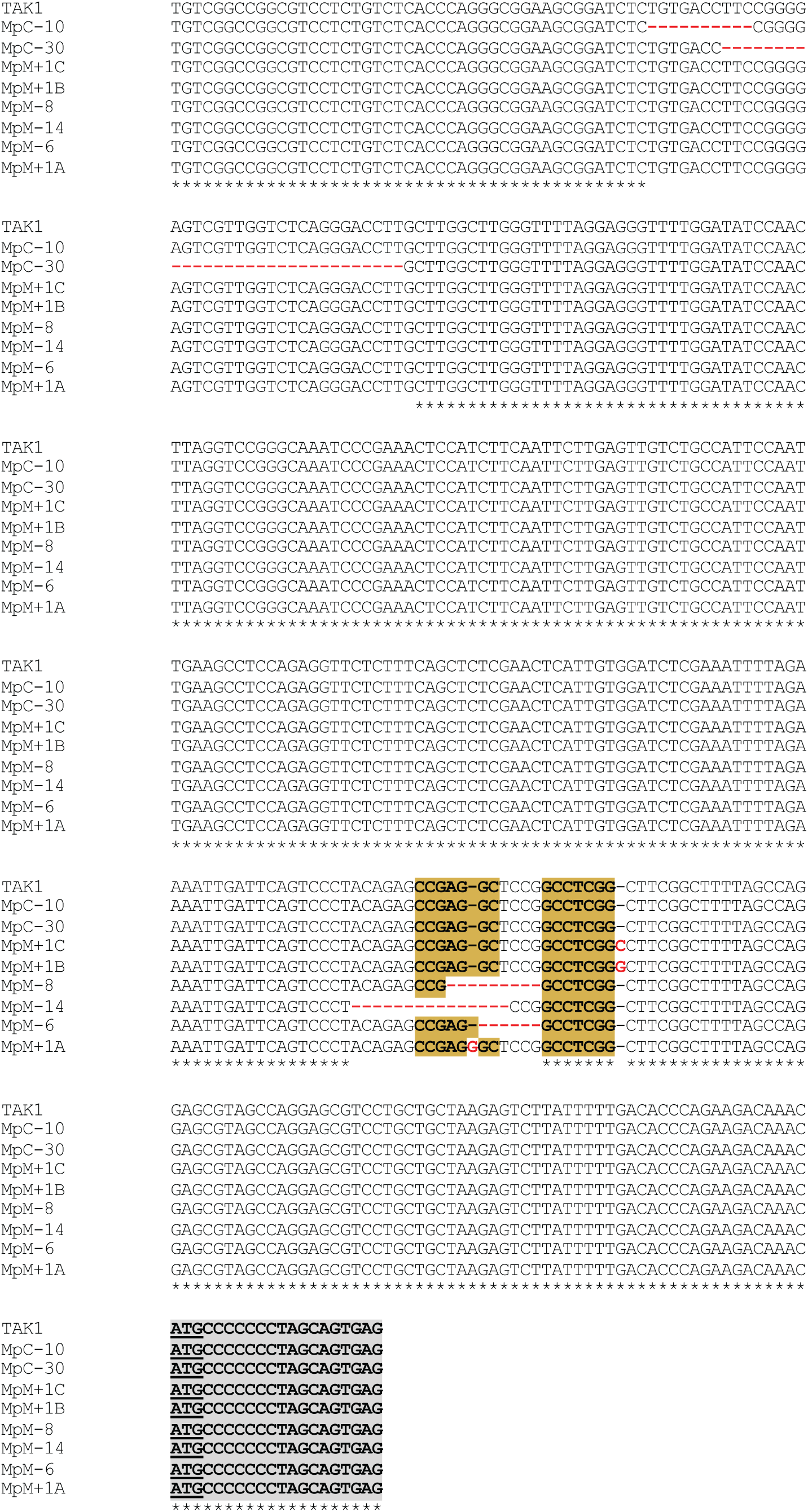
Sequence alignment of CRISPR–Cas9–induced *Marchantia adc-box* mutant alleles. Sequence alignment of the *Marchantia polymorpha* MpADC upstream region in WT (TAK1), upstream control mutants (MpC), and *adc-box* mutant lines (MpM). Deleted nucleotides are indicated by red dashes, and inserted nucleotides are shown in red letters. Conserved regions corresponding to the predicted GC2 and GC3 pairing elements of the RDRβ stem are highlighted in yellow background. Underlined codons represent start/stop codons of uORFs. Gray background indicates the start of the *MpADC* CDS. This alignment shows the positions and sizes of CRISPR-induced insertions and deletions used for functional analyses.

**Supplementary Figure 2.**
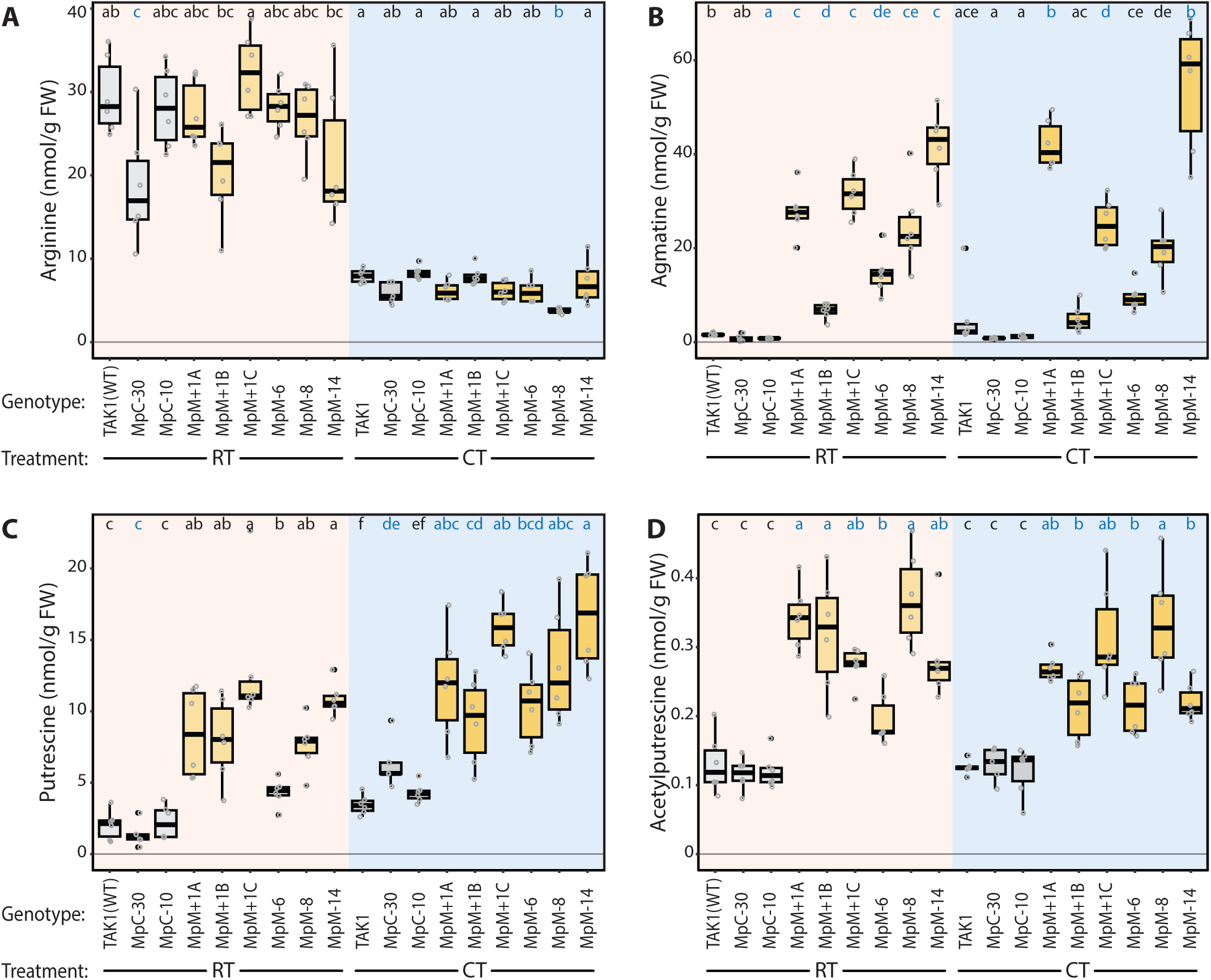
Disruption of the Marchantia ADC-box alters polyamine pathway metabolites. Quantification of polyamine pathway metabolites in WT (TAK1), upstream control mutants (MpC), and ADC-box mutants (MpM) under room temperature (RT) and cold treatment (CT) conditions. (A) Arginine, (B) agmatine, (C) putrescine, and (D) acetyl-putrescine levels determined by LC-MS and expressed as nmol/g fresh weight (FW).Boxplots show medians (center lines), interquartile ranges (boxes), and 1.5× interquartile ranges (whiskers), with individual biological replicates indicated as points. Different letters denote statistically significant differences (ANOVA followed by Tukey’s HSD, P < 0.05). These data complement Figure 1 by illustrating broader metabolic consequences of ADC-box disruption beyond ADC activity.

**Supplementary Figure 3.**
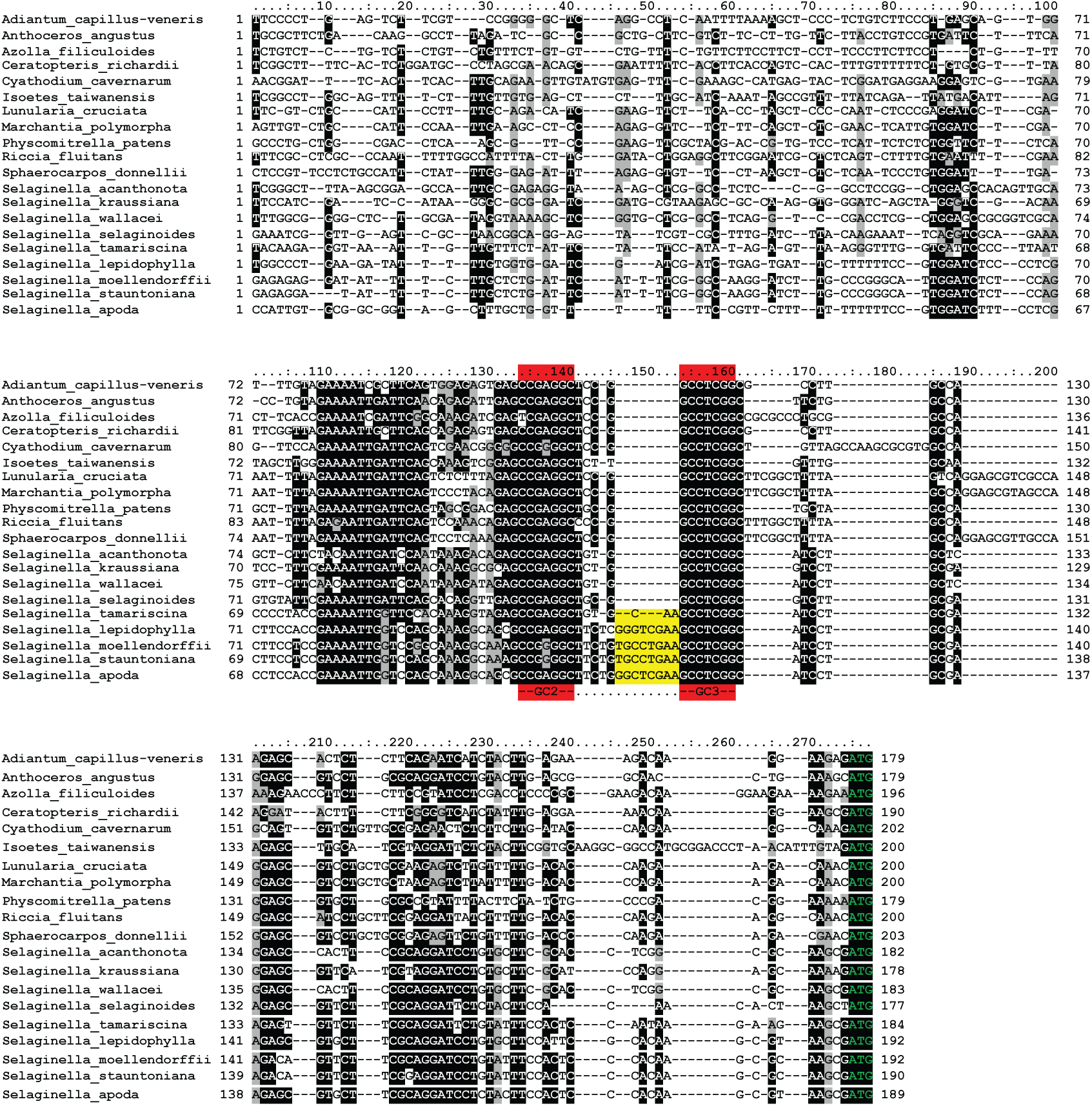
Conservation and variation of ADC-box sequences in seedless plants. Multiple sequence alignment of *ADC* upstream regions from representative seedless plant species, including bryophytes, ferns, and lycophytes. Conserved GC-rich regions corresponding to the predicted GC2 and GC3 elements are evident across species (red background colour), whereas sequence variation is primarily observed in flanking regions and in the loop between paired stems. Yellow background colour indicates expansion of the loop region in *Selaginella* species. This analysis supports the evolutionary conservation of the GC2/GC3 pairing region as the structural core of the ancestral ADC-box.

**Supplementary Figure 4.**
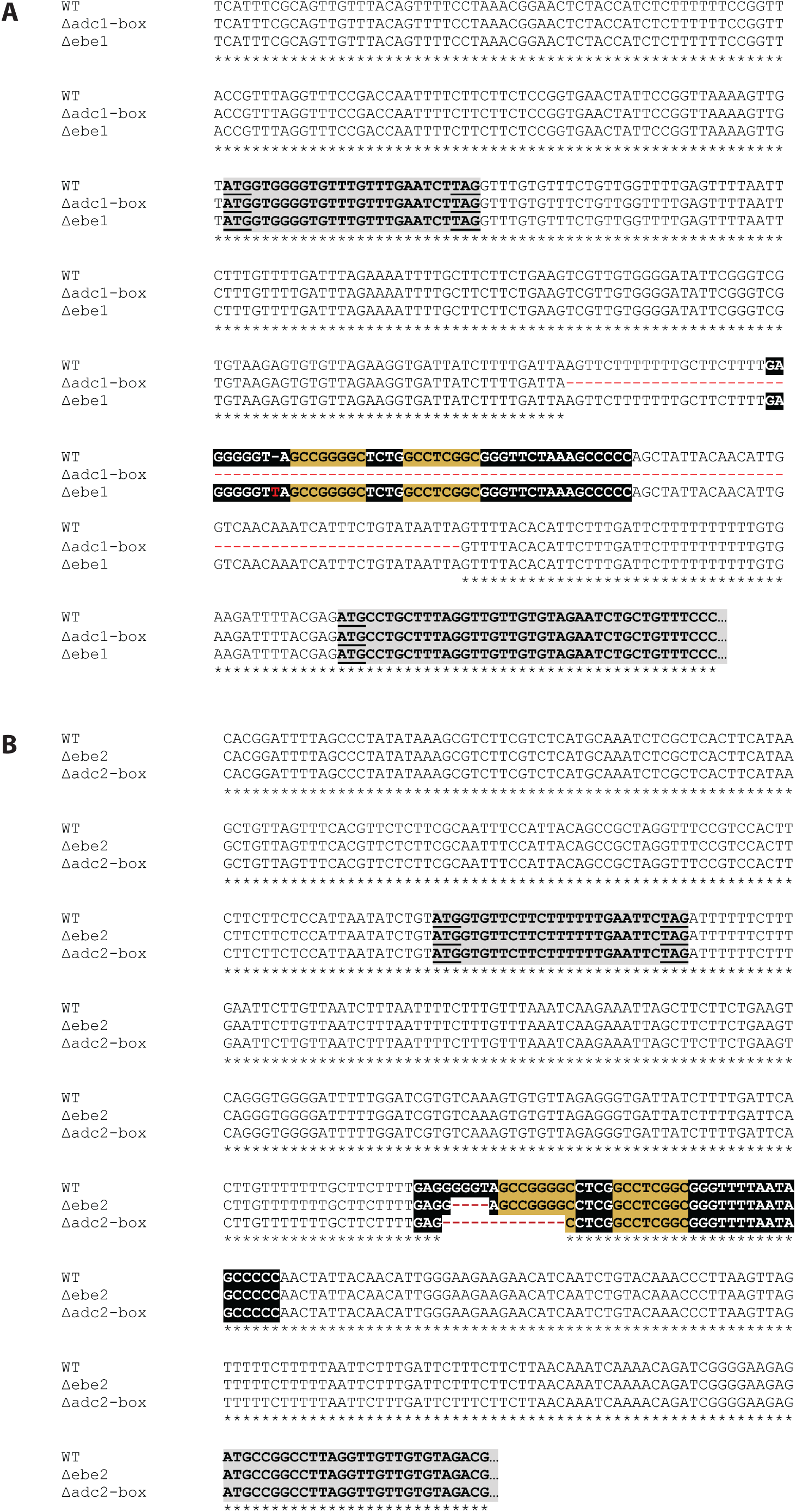
Alignment of CRISPR–Cas9-induced tomato ADC-box mutant alleles in *SlADC1* and *SlADC2* upstream regions. (A, B) Sequence alignments of the 5′ upstream regions of *SlADC1* (A) and *SlADC2* (B) from wild type (WT), effector binding element (*ebe*) mutants (*Δebe1, Δebe2*), and *adc-box* deletion mutants (*Δadc1-box*, *Δadc2-box*). Deleted nucleotides are indicated by red dashes, and insertions are shown in red font. Conserved GC-rich regions corresponding to the ADC-box stems (GC1–GC4) are highlighted in yellow. Underlined codons represent start/stop codons of uORFs. Small *ebe* mutations (*Δebe1*, *Δebe2*) retain the GC2/GC3 pairing region, whereas larger adc-box deletions (*Δadc1-box*, *Δadc2-box*) disrupt or remove this predicted structural core.

**Supplementary Figure 5.**
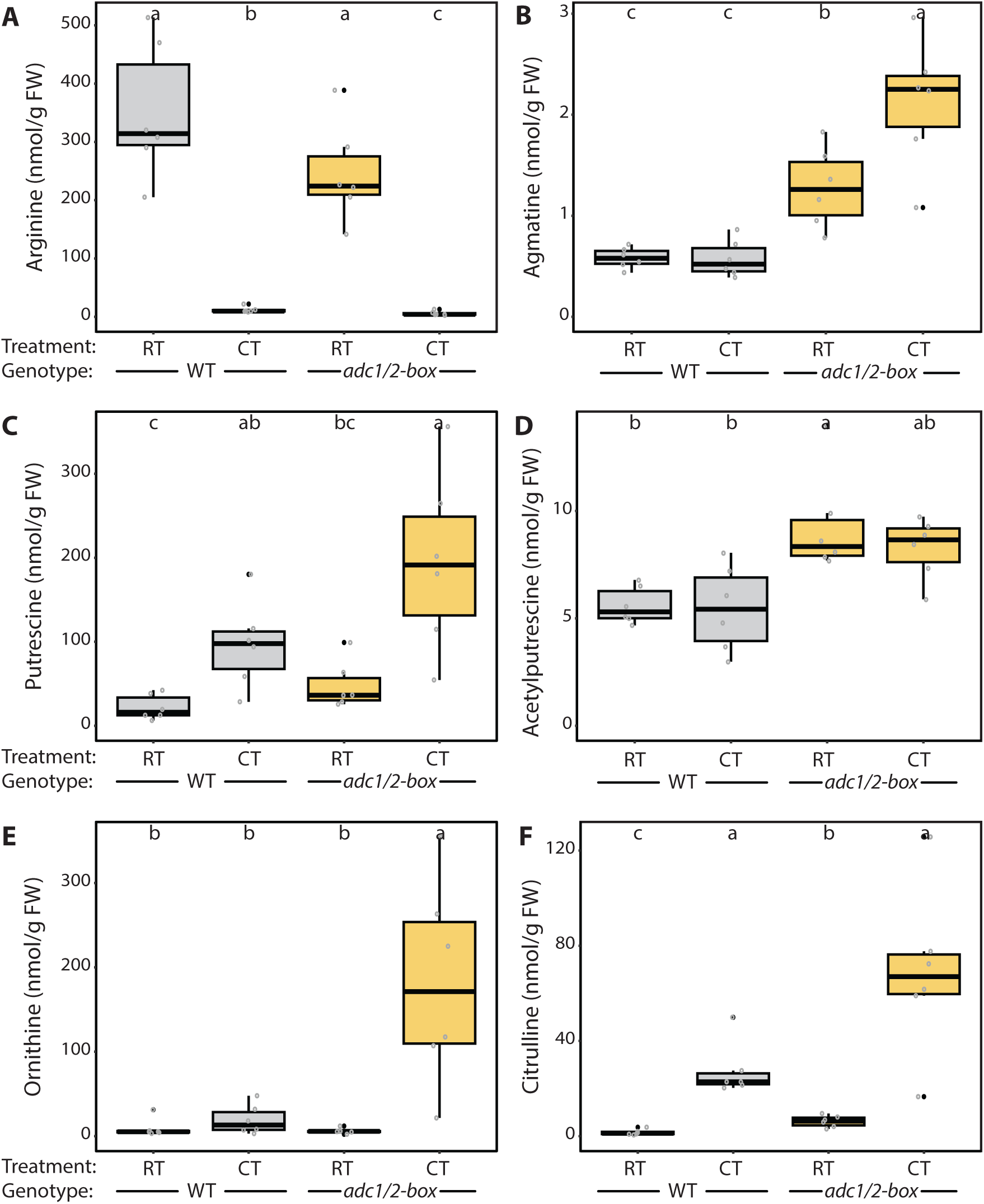
Disruption of the tomato *ADC-box* alters polyamine pathway metabolites. (A–F) Quantification of selected metabolites of the polyamine pathway and associated amino acid metabolism in wild type (WT; grey) and *Δadc1/2-box* mutant (yellow) tomato plants under room temperature (RT) and cold treatment (CT) conditions. Metabolites were measured by LC-MS and are expressed as nmol/g fresh weight (FW). (A) Arginine, (B) agmatine, (C) putrescine, (D) acetylputrescine, (E) ornithine, and (F) citrulline.Boxplots show medians (center lines), interquartile ranges (boxes), and 1.5× interquartile ranges (whiskers), with individual biological replicates indicated as points. Different letters denote statistically significant differences between groups (ANOVA followed by Tukey’s HSD, P < 0.05). Disruption of the *ADC-box* results in increased accumulation of metabolites proximal to ADC activity, particularly agmatine and putrescine, with additional effects on upstream and connected metabolites, indicating altered flux through the PA network.

**Supplementary Figure 6.**
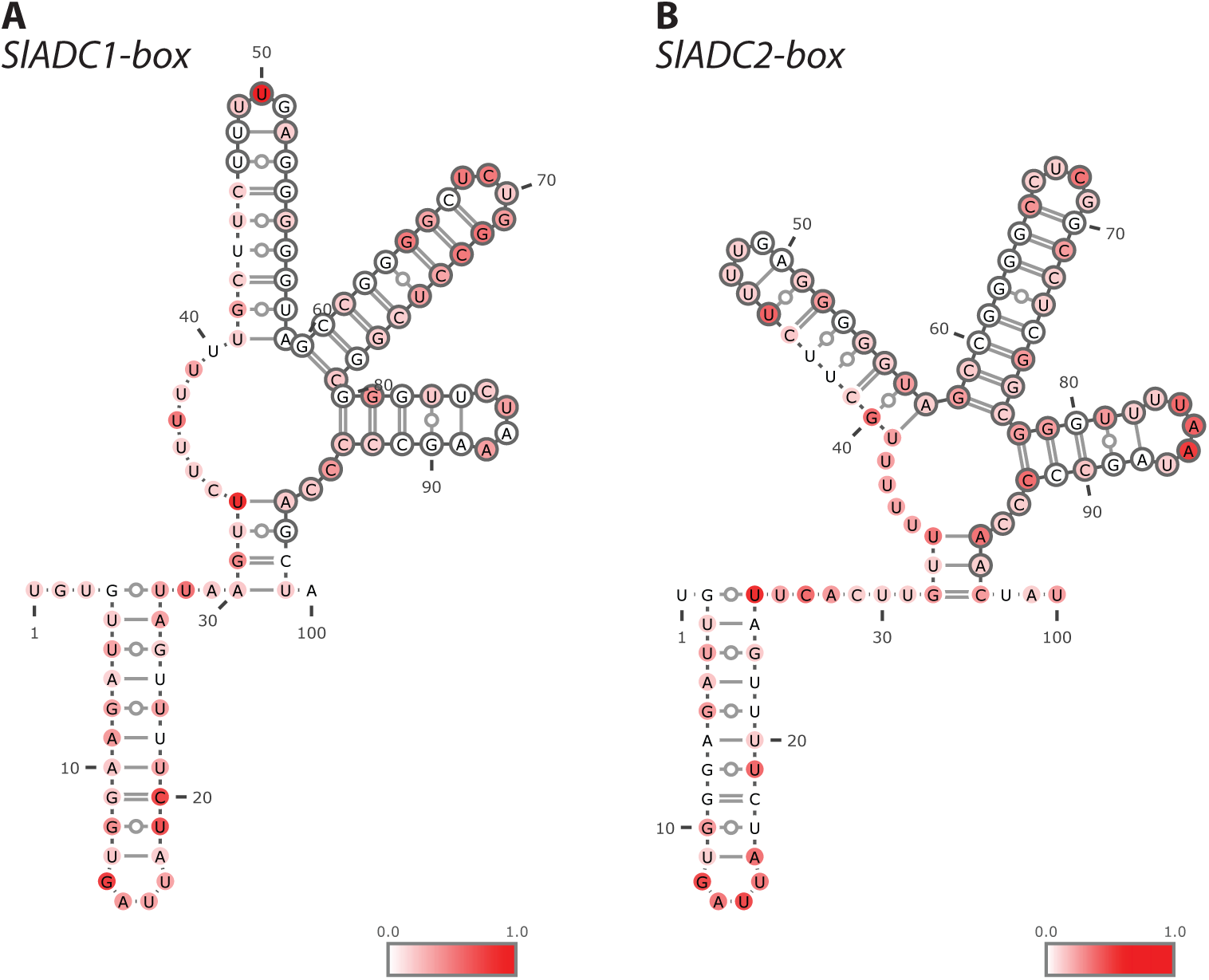
SHAPE-derived reactivity profiles of SlADC1- and SlADC2-box RNA secondary structures. (A, B) SHAPE-informed RNA secondary structure models of the *SlADC1-box* (A) and *SlADC2-box* (B). Nucleotides are coloured according to their SHAPE reactivity values, as indicated by the scale (0.0–1.0), with low reactivity (white) corresponding to structurally constrained or base-paired nucleotides and high reactivity (red) indicating flexible, single-stranded regions. Predicted base-pairing interactions are shown by connecting lines, and nucleotide positions are indicated. The SHAPE reactivity profiles support a three-stem architecture in both ADC-box elements, with low reactivity in stem regions and elevated reactivity in loop regions.

**Supplementary Figure 7.**
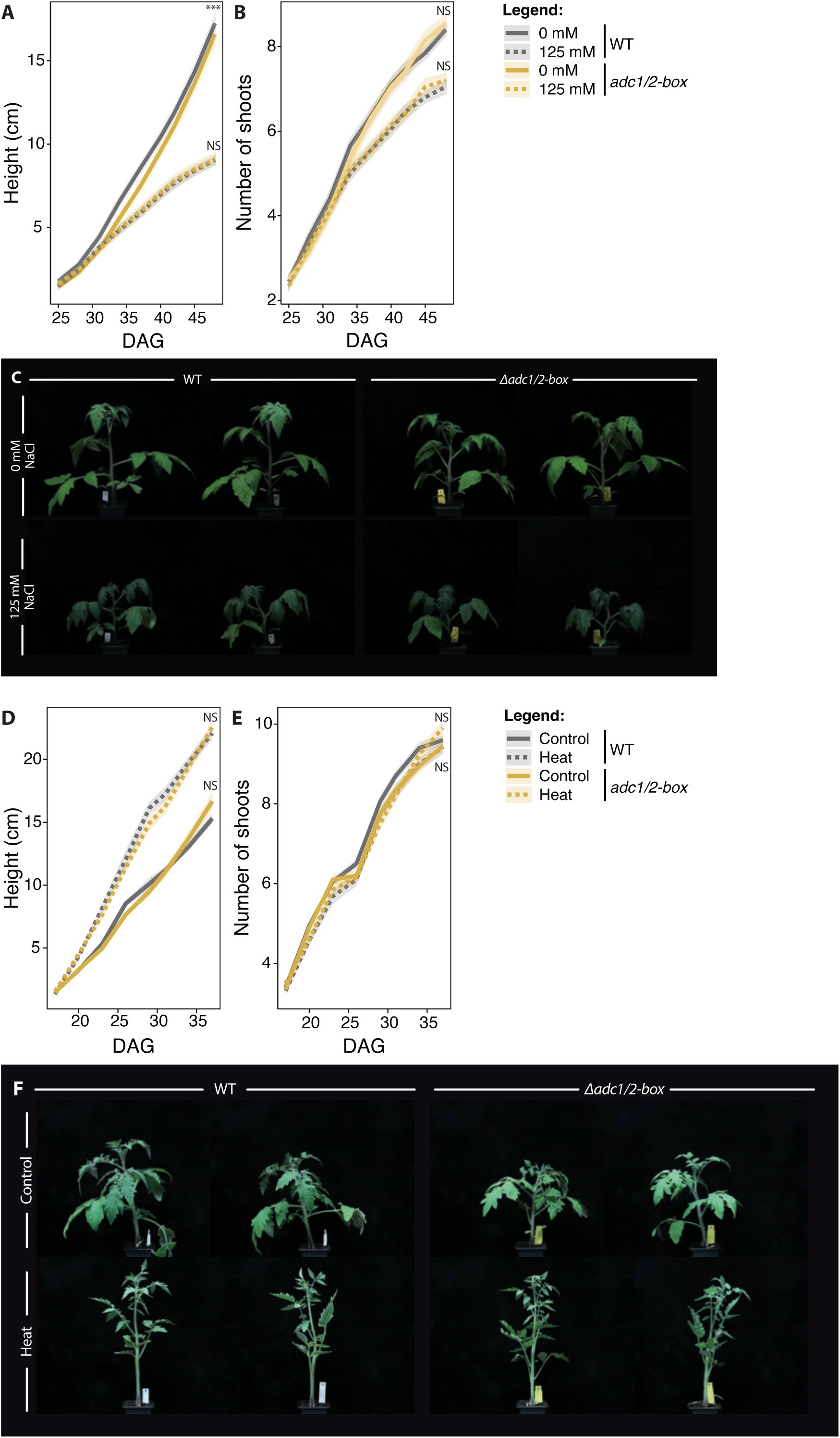
No detectable phenotypic di\erences between WT and Δadc1/2-box mutants under salt or heat stress. (A, B) Growth analysis of wild type (WT; grey) and *Δadc1/2-box* mutant (yellow) tomato plants subjected to salt stress (0 or 125 mM NaCl). Plant height (A) and number of shoots (B) were recorded over time (days after germination, DAG). (C) Representative images of WT and *Δadc1/2-box* plants grown under control (0 mM NaCl) or salt stress (125 mM NaCl) conditions. (D, E) Growth analysis of WT and *Δadc1/2-box* plants under control and heat stress conditions. Plant height (D) and number of shoots (E) were recorded over time (DAG). (F) Representative images of WT and *Δadc1/2-box* plants grown under control or heat stress conditions. Lines represent mean values ± SE. Statistical significance between genotypes within each treatment is indicated (NS, not significant; ***P < 0.001). Despite clear growth differences between control and stress conditions, no consistent or statistically significant differences were observed between WT and *Δadc1/2-box* plants under either salt or heat stress, indicating that disruption of the *ADC-box* does not measurably affect plant growth under these abiotic stress conditions.

**Supplementary Figure 8.**
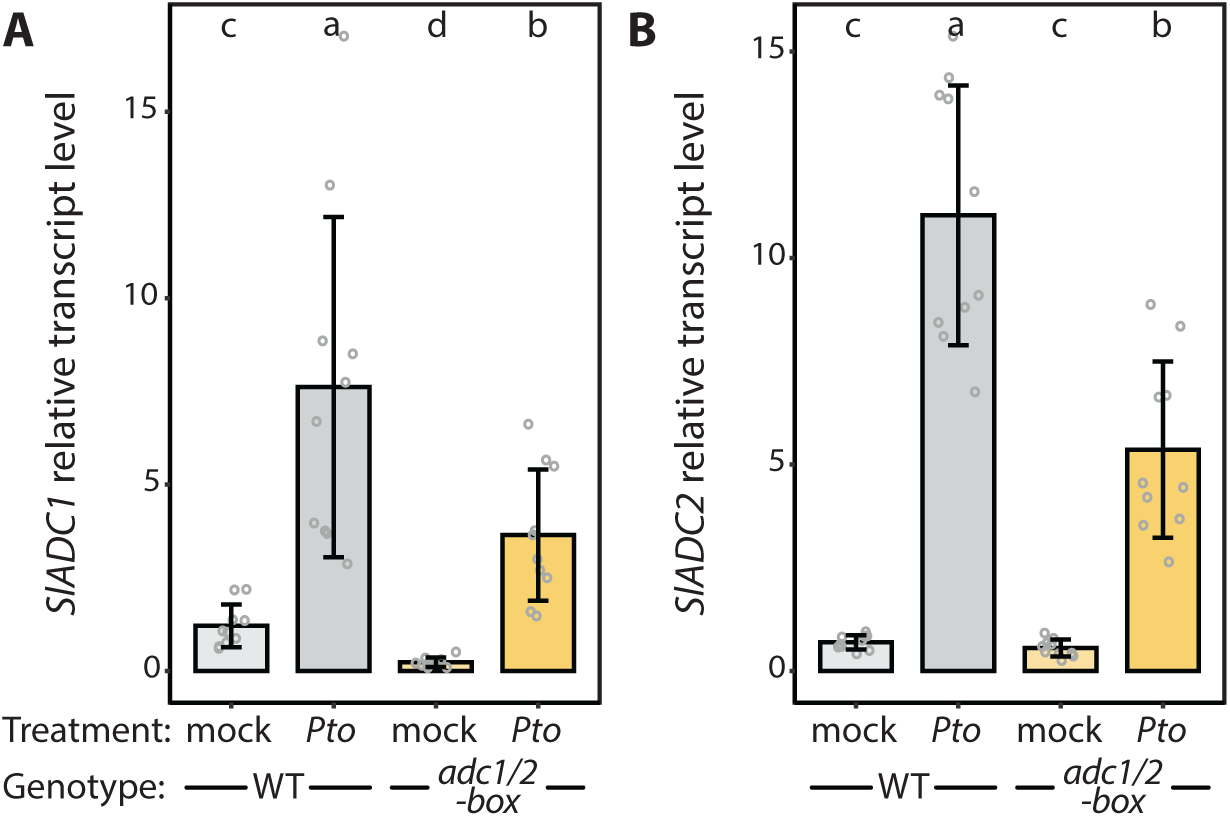
*Pto* infection induces SlADC1 and SlADC2 expression. (A, B) Relative transcript levels of *SlADC1 (A)* and *SlADC2 (B)* in wild type (WT; grey) and *Δadc1/2-box* mutant (yellow) tomato leaves following mock treatment or infection with *Pseudomonas syringae pv. tomato* (*Pto*). Transcript levels were determined by RT–qPCR and are shown relative to mock-treated WT. Bars represent mean values ± SE, with individual biological replicates indicated as points. Different letters denote statistically significant differences between groups (ANOVA followed by Tukey’s HSD, P < 0.05). *Pto* infection strongly induces *SlADC1* and *SlADC2* transcript accumulation in both WT and *Δadc1/2-box* plants, with no significant differences between genotypes under either mock or infected conditions, indicating that *ADC-box* mutations do not affect transcriptional induction of *SlADC* genes.

**Supplementary Table 1:**
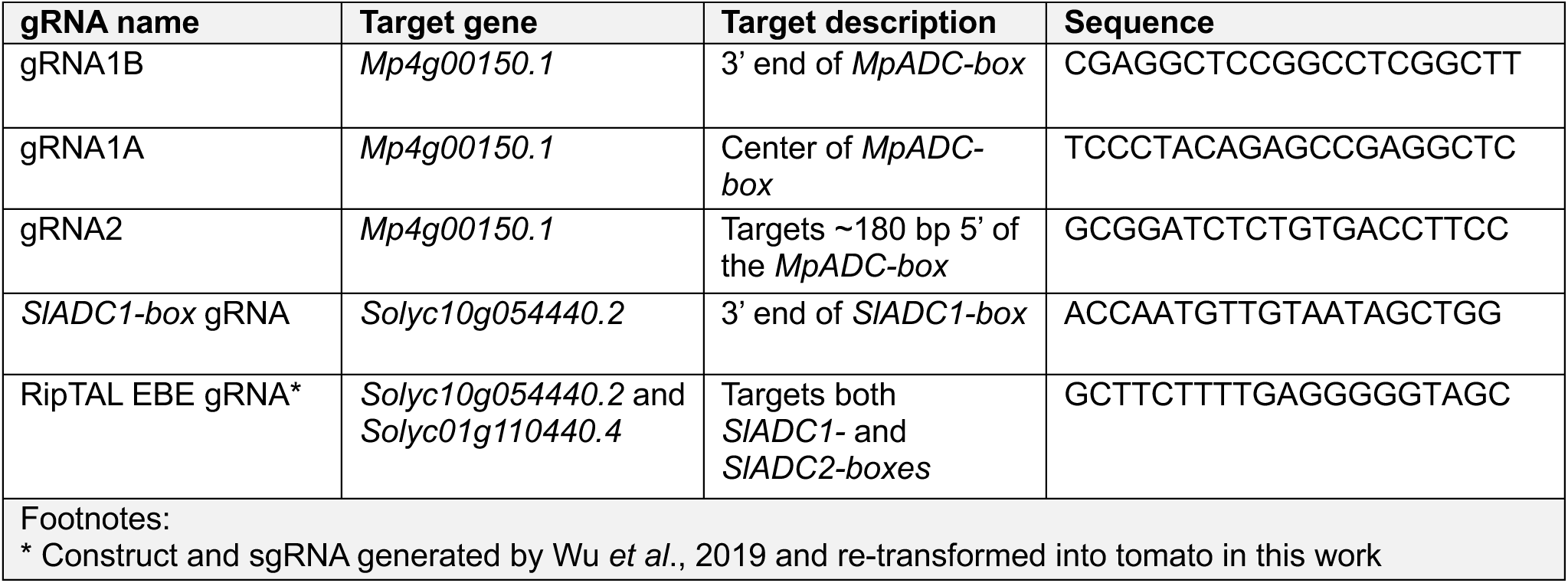
gRNAs used in this study.

**Supplementary Table 2:**
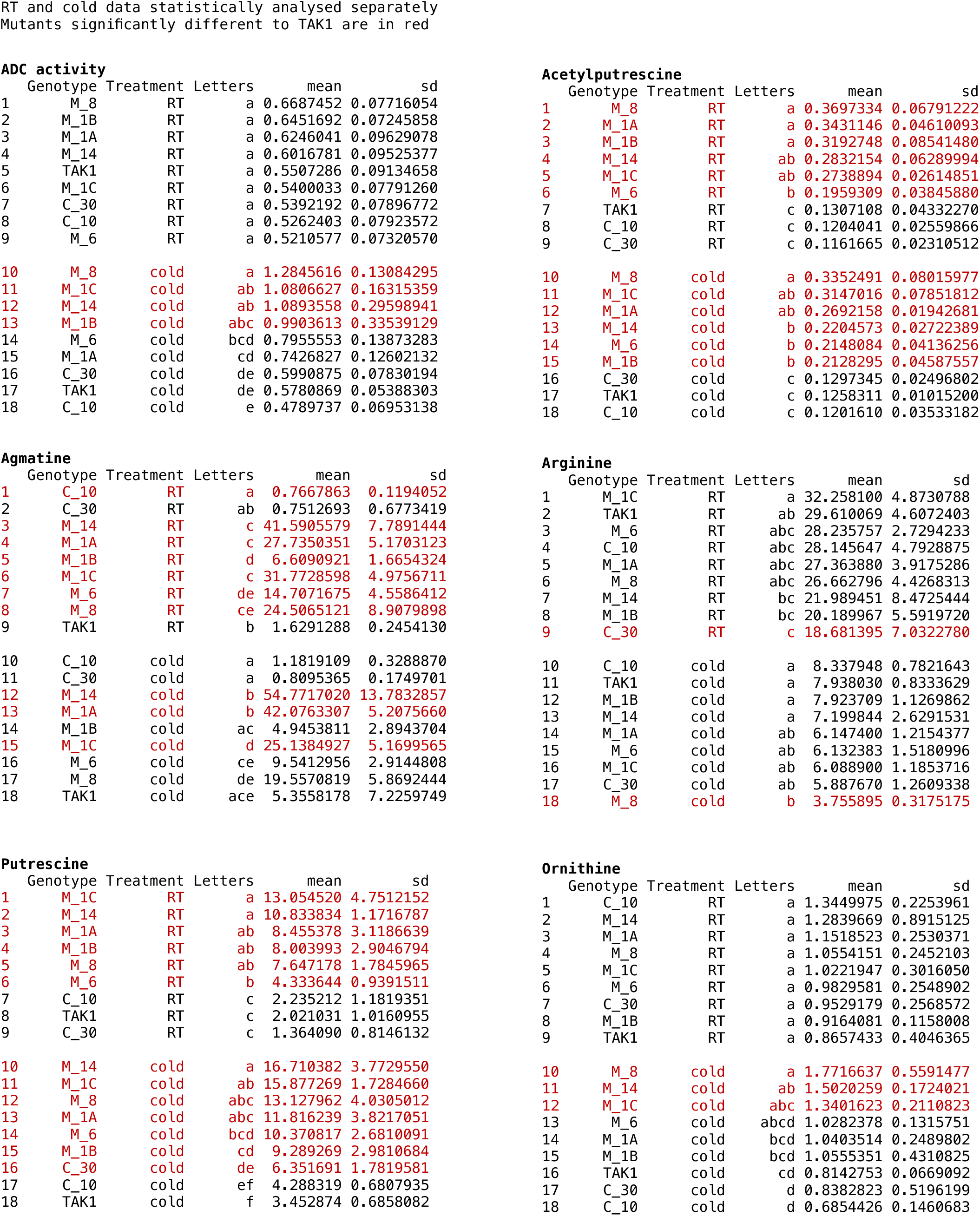
*M.polymorpha ADC-box* mutants, control lines, and TAK1 LC-MS quantified metabolite statistics.

**Supplementary Table 3:** Tomato *ADC-box* mutants and WT LC-MS quantified metabolite statistics.

ADC activity
|  | Sample.name | Letters | mean | sd |
| --- | --- | --- | --- | --- |
| 1 | *adc12box_Cold* | a | 129.212040 | 49.6064962 |
| 2 | WT_Cold | b | 13.950875 | 11.0712833 |
| 3 | *adc12box_RT* | c | 1.832230 | 0.2402413 |
| 4 | WT_RT | c | 1.178028 | 0.1182525 |

Agmatine
|  | Sample.name | Letters | mean | sd |
| --- | --- | --- | --- | --- |
| 1 | *adc12box_Cold* | a | 2.1215572 | 0.6391763 |
| 2 | *adc12box_RT* | b | 1.2791525 | 0.3934407 |
| 3 | WT_RT | c | 0.5820749 | 0.1024683 |
| 4 | WT_Cold | c | 0.5754929 | 0.1820674 |

Putrescine
|  | Sample.name | Letters | mean | sd |
| --- | --- | --- | --- | --- |
| 1 | *adc12box_Cold* | a | 195.44314 | 106.94823 |
| 2 | WT_Cold | ab | 96.48365 | 51.87886 |
| 3 | *adc12box_RT* | bc | 48.25271 | 28.29779 |
| 4 | WT_RT | c | 21.91443 | 14.90924 |

Arginine
|  | Sample.name | Letters | mean | sd |
| --- | --- | --- | --- | --- |
| 1 | WT_RT | a | 351.175135 | 116.876522 |
| 2 | *adc12box_RT* | a | 245.877072 | 84.742527 |
| 3 | WT_Cold | b | 11.754305 | 5.096255 |
| 4 | *adc12box_Cold* | c | 5.707633 | 3.770123 |

Ornithine
|  | Sample.name | Letters | mean | sd |
| --- | --- | --- | --- | --- |
| 1 | *adc12box_Cold* | a | 181.523678 | 121.451849 |
| 2 | WT_Cold | b | 19.356920 | 17.386783 |
| 3 | WT_RT | b | 9.241284 | 10.841858 |
| 4 | *adc12box_RT* | b | 6.025424 | 3.194953 |

Acetylputrescine
|  | Sample.name | Letters | mean | sd |
| --- | --- | --- | --- | --- |
| 1 | *adc12box_RT* | a | 9.343671 | 2.3820459 |
| 2 | *adc12box_Cold* | ab | 8.259754 | 1.4214921 |
| 3 | WT_RT | b | 5.593026 | 0.8672813 |
| 4 | WT_Cold | b | 5.453921 | 1.9922310 |

Citrulline
|  | Sample.name | Letters | mean | sd |
| --- | --- | --- | --- | --- |
| 1 | *adc12box_Cold* | a | 68.851301 | 35.234617 |
| 2 | WT_Cold | a | 27.467690 | 11.274992 |
| 3 | *adc12box_RT* | b | 6.203772 | 2.430837 |
| 4 | WT_RT | c | 1.560582 | 1.176639 |

**Supplementary Table 4:**
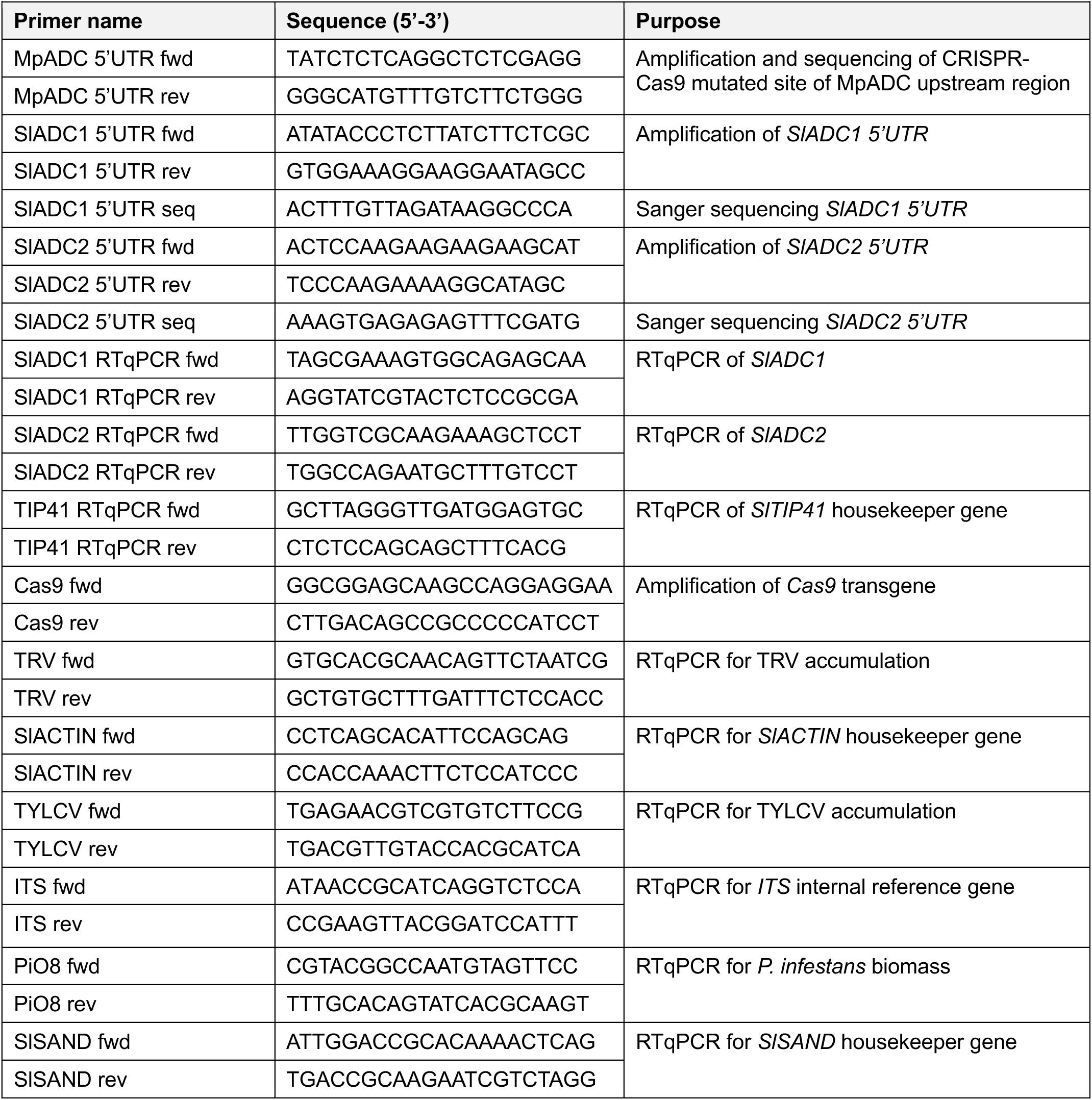
Primers used in this study.

## References

Alcázar R, García-Martínez JL, Cuevas JC, Tiburcio AF, Altabella T (2005) Overexpression of *ADC2* in *Arabidopsis* induces dwarfism and late-flowering through GA deficiency. Plant Journal 43: 425–436

Bassard JE, Ullmann P, Bernier F, Werck-Reichhart D (2010) Phenolamides: bridging polyamines to the phenolic metabolism. Phytochemistry 71: 1808–1824

Blázquez MA (2024) Polyamines: their role in plant development and stress. Annual Review of Plant Biology 75: 95–117

Chen D, Shao Q, Yin L, Younis A, Zheng B (2019) Polyamine function in plants: metabolism, regulation on development, and roles in abiotic stress responses. Frontiers in Plant Science 9: 1945

Fernandez-Calvino L, Osorio S, Hernandez ML, Hamada IB, Del Toro FJ, Donaire L, Yu A, Bustos R, Fernie AR, Martinez-Rivas JM, Llave C (2014) Virus-induced alterations in primary metabolism modulate susceptibility to Tobacco rattle virus in *Arabidopsis*. Plant Physiology 166: 1821–1838

Fry W (2008) *Phytophthora infestans*: the plant (and R gene) destroyer. Molecular Plant Pathology 9: 385–402

Gallas N, Li XX, von Roepenack-Lahaye E, Schandry N, Jiang YX, Wu DS, Lahaye T (2024) An ancient *cis*-element targeted by *Ralstonia solanacearum* TALE-like effectors facilitates the development of a promoter trap that could confer broad-spectrum wilt resistance. Plant Biotechnology Journal 22: 602–616

Gerlin L, Baroukh C, Genin S (2021) Polyamines: double agents in disease and plant immunity. Trends in Plant Science 26: 1061–1071

Giovannetti M, Mosse B (1980) An evaluation of techniques for measuring vesicular arbuscular mycorrhizal infection in roots. New Phytologist 84: 489–500

Gonzalez ME, Jasso-Robles FI, Flores-Hernandez E, Rodriguez-Kessler M, Pieckenstain FL (2021) Current status and perspectives on the role of polyamines in plant immunity. Annals of Applied Biology 178: 244–255

Jacobs TB, LaFayette PR, Schmitz RJ, Parrott WA (2015) Targeted genome modifications in soybean with CRISPR/Cas9. BMC Biotechnology 15: 16

Jimenez-Bremont JF, Chavez-Martinez AI, Ortega-Amaro MA, Guerrero-Gonzalez ML, Jasso-Robles FI, Maruri-Lopez I, Liu JH, Gill SS, Rodriguez-Kessler M (2022) Translational and post-translational regulation of polyamine metabolic enzymes in plants. Journal of Biotechnology 344: 1–10

Jiménez-Bremont JF, Marina M, Guerrero-González ML, Rossi FR, Sánchez-Rangel D, Rodríguez-Kessler M, Ruiz O, Gárriz A (2014) Physiological and molecular implications of plant polyamine metabolism during biotic interactions. Frontiers in Plant Science 5: 95

Judelson HS, Randall TA (1998) Families of repeated DNA in the oomycete *Phytophthora infestans* and their distribution within the genus. Genome 41: 605–615

Kim SH, Kim SH, Yoo SJ, Min KH, Nam SH, Cho BH, Yang KY (2013) Putrescine regulating by stress-responsive MAPK cascade contributes to bacterial pathogen defense in *Arabidopsis*. Biochemical and Biophysical Research Communications 437: 502–508

Kovach ME, Elzer PH, Hill DS, Robertson GT, Farris MA, Roop RM II, Peterson KM (1995) Four new derivatives of the broad-host-range cloning vector pBBR1MCS, carrying different antibiotic-resistance cassettes. Gene 166: 175–176

Li T, Huang Y, Xu ZS, Wang F, Xiong AS (2019) Salicylic acid-induced differential resistance to the Tomato yellow leaf curl virus among resistant and susceptible tomato cultivars. BMC Plant Biology 19: 173

Liu C, Atanasov KE, Arafaty N, Murillo E, Tiburcio AF, Zeier J, Alcazar R (2020) Putrescine elicits ROS-dependent activation of the salicylic acid pathway in *Arabidopsis thaliana*. Plant, Cell & Environment 43: 2755–2768

Liu Y, Schiff M, Marathe R, Dinesh-Kumar SP (2002) Tobacco Rar1, EDS1 and NPR1/NIM1-like genes are required for N-mediated resistance to tobacco mosaic virus. Plant Journal 30: 415–429

Lowe-Power TM, Hendrich CG, von Roepenack-Lahaye E, Li B, Wu D, Mitra R, Dalsing BL, Ricca P, Naidoo J, Cook D, Jancewicz A, Masson P, Thomma B, Lahaye T, Michael AJ, Allen C (2018) Metabolomics of tomato xylem sap during bacterial wilt reveals *Ralstonia solanacearum* produces abundant putrescine, a metabolite that accelerates wilt disease. Environmental Microbiology 20: 1330–1349

Lozano-Durán R (2024) Viral recognition and evasion in plants. Annual Review of Plant Biology 75: 655–677

Macho AP, Guidot A, Barberis P, Beuzon CR, Genin S (2010) A competitive index assay identifies several *Ralstonia solanacearum* type III effector mutant strains with reduced fitness in host plants. Molecular Plant-Microbe Interactions 23: 1197–1205

Mason G, Caciagli P, Accotto GP, Noris E (2008) Real-time PCR for the quantitation of Tomato yellow leaf curl Sardinia virus in tomato plants and in *Bemisia tabaci*. Journal of Virological Methods 147: 282–289

Medina-Puche L, Tan H, Dogra V, Wu M, Rosas-Diaz T, Wang L, Ding X, Zhang D, Fu X, Kim C, Lozano-Durán R (2020) A defense pathway linking plasma membrane and chloroplasts and co-opted by pathogens. Cell 182: 1109–1124.e25

Michael AJ (2016) Biosynthesis of polyamines and polyamine-containing molecules. Biochemical Journal 473: 2315–2329

Morcillo RJL, Zhao A, Tamayo-Navarrete MI, Garcia-Garrido JM, Macho AP (2020) Tomato root transformation followed by inoculation with *Ralstonia solanacearum* for straightforward genetic analysis of bacterial wilt disease. Journal of Visualized Experiments 160: e60302

Murillo E, Martínez-Seidel F, Atanasov KE, Gentry-Torfer D, Firmino AAP, Erban A, Nie S, Leeming MG, Suwanchaikasem P, Boughton BA, Williamson NA, Roessner U, Kopka J, Alcázar R (2025) Polyamines and flg22 reshape the ribosomal protein composition of actively translating ribosomes in plants. Plant Physiology and Biochemistry 220: 109684

Murray MG, Thompson WF (1980) Rapid isolation of high molecular weight plant DNA. Nucleic Acids Research 8: 4321–4325

Naito Y, Hino K, Bono H, Ui-Tei K (2015) CRISPRdirect: software for designing CRISPR/Cas guide RNA with reduced oh-target sites. Bioinformatics 31: 1120–1123

Nakagawa T, Suzuki T, Murata S, Nakamura S, Hino T, Maeo K, Tabata R, Kawai T, Tanaka K, Niwa Y, Watanabe Y, Nakamura K, Kimura T, Ishiguro S (2007) Improved Gateway binary vectors: high-performance vectors for creation of fusion constructs in transgenic analysis of plants. Bioscience, Biotechnology, and Biochemistry 71: 2095–2100

Oñate-Sánchez L, Vicente-Carbajosa J (2008) DNA-free RNA isolation protocols for *Arabidopsis thaliana*, including seeds and siliques. BMC Research Notes 1: 93

Pathak MR, Teixeira da Silva JA, Wani SH (2014) Polyamines in response to abiotic stress tolerance through transgenic approaches. GM Crops & Food 5: 87–96

Ritchie ES, von Roepenack-Lahaye E, Perrett D, Wu D, Lahaye T (2025) Optimized LC-MS method for simultaneous polyamine profiling and ADC/ODC activity quantification and evidence that ADCs are indispensable for flower development in tomato. Frontiers in Plant Science 16: 1636076

Sánchez-Rangel D, Chávez-Martinez AI, Rodríguez-Hernández AA, Maruri-López I, Urano K, Shinozaki K, Jiménez-Bremont JF (2016) Simultaneous silencing of two arginine decarboxylase genes alters development in *Arabidopsis*. Frontiers in Plant Science 7: 300

Sharma K, Gupta S, Thokchom SD, Jangir P, Kapoor R (2021) Arbuscular mycorrhiza-mediated regulation of polyamines and aquaporins during abiotic stress: deep insights on the recondite players. Frontiers in Plant Science 12: 642101

Si-Ammour A, Mauch-Mani B, Mauch F (2003) Quantification of induced resistance against *Phytophthora* species expressing GFP as a vital marker: β-aminobutyric acid but not BTH protects potato and *Arabidopsis* from infection. Molecular Plant Pathology 4: 237–248

Spitale RC, Crisalli P, Flynn RA, Torre EA, Kool ET, Chang HY (2013) RNA SHAPE analysis in living cells. Nature Chemical Biology 9: 18–20

Sugano SS, Shirakawa M, Takagi J, Matsuda Y, Shimada T, Hara-Nishimura I, Kohchi T (2014) CRISPR/Cas9-mediated targeted mutagenesis in the liverwort *Marchantia polymorpha* L. Plant and Cell Physiology 55: 475–481

Tan QW, Lim PK, Chen Z, Pasha A, Provart N, Arend M, Nikoloski Z, Mutwil M (2023) Cross-stress gene expression atlas of *Marchantia polymorpha* reveals the hierarchy and regulatory principles of abiotic stress responses. Nature Communications 14: 986

Tanizawa Y, Mochizuki T, Yagura M, Sakamoto M, Fujisawa T, Kawamura S, Shimokawa E, Yamaoka S, Nishihama R, Bowman JL, Berger F, Yamato KT, Kohchi T, Nakamura Y (2026) MarpolBase: genome database for *Marchantia polymorpha* featuring high-quality reference genome sequences. Plant and Cell Physiology 67: 377–388

Torabi S, Varshney K, Villaécija-Aguilar JA, Keymer A, Gutjahr C (2021) Controlled assays for phenotyping the effects of strigolactone-like molecules on arbuscular mycorrhiza development. Methods in Molecular Biology 2309: 157–177

Upadhyay RK, Fatima T, Handa AK, Mattoo AK (2020) Polyamines and their biosynthesis/catabolism genes are differentially modulated in response to heat versus cold stress in tomato leaves (*Solanum lycopersicum* L.). Cells 9: 1749

Urano K, Hobo T, Shinozaki K (2005) *Arabidopsis* ADC genes involved in polyamine biosynthesis are essential for seed development. FEBS Letters 579: 1557–1564

Vilas JM, Romero FM, Rossi FR, Marina M, Maiale SJ, Calzadilla PI, Pieckenstain FL, Ruiz OA, Garriz A (2018) Modulation of plant and bacterial polyamine metabolism during the compatible interaction between tomato and *Pseudomonas syringae*. Journal of Plant Physiology 231: 281–290

Wang L, Tan H, Wu M, Jimenez-Gongora T, Tan L, Lozano-Durán R (2017) Dynamic virus-dependent subnuclear localization of the capsid protein from a geminivirus. Frontiers in Plant Science 8: 2165

Wang W, Paschalidis K, Feng JC, Song J, Liu JH (2019) Polyamine catabolism in plants: a universal process with diverse functions. Frontiers in Plant Science 10: 561

Wittmann J, Brancato C, Berendzen KW, Dreiseikelmann B (2016) Development of a tomato plant resistant to *Clavibacter michiganensis* using the endolysin gene of bacteriophage CMP1 as a transgene. Plant Pathology 65: 496–502

Wu D, von Roepenack-Lahaye E, Buntru M, de Lange O, Schandry N, Pérez-Quintero AL, Weinberg Z, Lowe-Power TM, Szurek B, Michael AJ, Allen C, Schillberg S, Lahaye T (2019) A plant pathogen type III effector protein subverts translational regulation to boost host polyamine levels. Cell Host & Microbe 26: 638–649.e5

Xin XF, He SY (2013) *Pseudomonas syringae* pv. tomato DC3000: a model pathogen for probing disease susceptibility and hormone signaling in plants. Annual Review of Phytopathology 51: 473–498

Xue H, Lozano-Durán R, Macho AP (2020) Insights into the root invasion by the plant pathogenic bacterium *Ralstonia solanacearum*. Plants 9: 516

Yang X, Qin H, Zhou Y, Mai Z, Chai X, Guo J, Kang Y, Zhong M (2025) HB52-PUT2 module-mediated polyamine shoot-to-root movement regulates salt stress tolerance in tomato. *Plant*, Cell & Environment 48: 5148–5163

Yang XF, Cheema J, Zhang YY, Deng HJ, Duncan S, Umar MI, Zhao JY, Liu Q, Cao XF, Kwok CK, Ding YL (2020) RNA G-quadruplex structures exist and function in vivo in plants. Genome Biology 21: 226

Yang K, Yan Q, Wang Y, Li S, Zhang M, Liu Z, Zhao L, Chen (2022) GmPAO-mediated polyamine catabolism enhances soybean *Phytophthora* resistance without growth penalty. Phytopathol Res 4, 35.

Zou YN, Zhang F, Srivastava AK, Wu QS, Kuca K (2020) Arbuscular mycorrhizal fungi regulate polyamine homeostasis in roots of trifoliate orange for improved adaptation to soil moisture deficit stress. Frontiers in Plant Science 11: 600792

